# IL-21 selectively augments cytotoxic potential of antigen-activated MAIT cells

**DOI:** 10.1101/2025.03.20.644465

**Authors:** Laura E. Wedlock, Rajesh Lamichhane, Selena C. L. Gilmer, Andrea J. Vernall, James E. Ussher

## Abstract

Mucosal-associated invariant T (MAIT) cells are unconventional cytotoxic T cells restricted by MHC class 1 related molecule, MR1. They are activated through their TCR by derivatives from microbial riboflavin synthesis or independently of TCR signalling via IL-12 and IL-18. Upon activation, MAIT cells upregulate cytotoxic molecules GrzB and perforin and lyse bacterially-infected cells. While cytokines act as co-stimulatory molecules that enhance MAIT cell activation, their specific role in regulating MAIT cell cytotoxicity remains unresolved. We show that the cytokine IL-21 enhances expression of GrzB and perforin on TCR or IL-12/IL-18-activated MAIT cells but has limited effect on MAIT cell cytokine production. Using a flow cytometry-based cytotoxic assay, we show priming MAIT cells with IL-21 enhances their ability to kill 5-OP-RU-treated B cell lines. We demonstrate a previously unexplored co-stimulatory role for IL-21 that selectively augments the cytotoxic potential of MAIT cells.

## Introduction

Mucosal-associated invariant T (MAIT) cells are an abundant unconventional T cell population enriched in mucosal sites, such as gut, lung, female genital mucosa, and oral mucosa (1-4) and represent up to 20-50% of total T cells in the liver (5, 6) and 1-10% of T cells in peripheral blood (PB).(7, 8) MAIT cells are characterised by expression of an ab T cell receptor (TCR), comprised of a semi-invariant α-chain Vα7.2-Jα33/12/20 associated with a limited repertoire of β chains, most commonly Vβ2/Vβ13 in humans.(2, 9-11) Restricted by major histocompatibility complex (MHC)-class 1 related molecule (MR1), the MAIT cell TCR recognises conserved pyrimidine derivatives from microbial riboflavin synthesis, namely 5-(2-oxopropylideneamino)-6-d-ribitylaminouracil (5-OP-RU).(12, 13)

Activated MAIT cells upregulate interferon (IFN)-γ, tumour-necrosis factor (TNF)-α, and/or interleukin (IL)-17A and produce cytotoxic molecules to lyse bacterially-infected cells.(14-19) Resting MAIT cells express high levels of granzyme (Grz) A, GrzK, and GrzM, but low or negligible levels of GrzB, perforin, and granulysin.(17, 19-22) GrzB and perforin are significantly upregulated in activated MAIT cells,(19) and serve as indicators of MAIT cell cytotoxic potential. Changes in MAIT cell cytotoxic granule content, induced by cytokines and/or *E. coli*, enhances TCR-mediated killing.(19) However, stimulation with anti-CD3/CD28 or by MR1 ligands alone, without these co-stimulatory signals, is not sufficient to trigger the full effector potential of MAIT cells.(23-25)

Cytokines can provide co-stimulation to regulate MAIT cell responses. A combination of IL-12 and IL-18 triggers TCR-independent activation of MAIT cells,(26) while cytokines IL-7 and IL-15 enhance MAIT cell cytotoxicity, IFN-γ production, and augment their response to bacteria.(22, 27, 28) Type 1 interferon (T1-IFN) signalling also modulates MAIT cell activation. IFN-α enhances MAIT cell response to 5-OP-RU,(29) anti-CD3,(30) and IL-12 and IL-18,(31) but alone induces only modest activation.(29) Thus, when combined with IL-12/IL-18 or TCR signalling, cytokines boost the activation and cytotoxic response of MAIT cells.

IL-21 is a pleiotropic type 1 cytokine mainly produced by activated CD4^+^ T cells (particularly T follicular helper (Tfh) and T helper 17 (Th17) cells)(32) and iNKT cells,(33) but is produced in smaller amounts by CD8^+^ T cells and MAIT cells in some contexts.(34-38) IL-21 predominantly signals through Janus kinase (JAK)–signal transducer and activator of transcription (STAT) pathway via heterodimers of IL-21R with the common gamma chain (γc) receptor.(32, 38-40) In T cells, IL-21 induces strong and sustained phosphorylation of STAT3, and to lesser degrees, STAT1, STAT5a, and STAT5b.(38) A previous study reported MAIT cells express IL-21R and IL-21 activates STAT3 in MAIT cells.(41) Additionally, individuals with loss-of-function mutations in IL-21R or STAT3, have reduced frequency of circulating MAIT and iNKT cells, and residual STAT3-deficient MAIT cells exhibit diminished IL-17A production.(41)

IL-21 enhances cytotoxicity of various immune cells, including neutrophils,(42) NK cells,(43) and CD8^+^ T cells,(44) however, direct function of IL-21 in modulating MAIT cell cytotoxicity has not been demonstrated. In the current study, we show that IL-21 enhances production of cytotoxic molecules by MAIT cells when associated with a TCR or cytokine signal, but has limited effect on TCR-mediated cytokine production. We demonstrate that IL-21 selectively augments the cytotoxic potential of human PB MAIT cells.

## Materials and Methods

### Cell isolation and culture

Human peripheral blood mononuclear cells (PBMCs) were isolated from Leukoreduction System Chambers (obtained from NZ Blood) using Lymphoprep™ (Axis-Shield, Dundee, UK) density gradient centrifugation. Isolated PBMCs were rested overnight and used for Vα7.2^+^ cell isolation the following day or frozen and stored at -80°C or in liquid nitrogen. For each experiment, frozen PBMCs were thawed, washed, and rested overnight in R10 (RPMI 1640 with L-glutamine (Life Technologies, Carlsbard, CA) and 10% FBS (Gibco™, NZ) with 100 μg/mL streptomycin + 100 units/mL penicillin (Sigma-Aldrich, St Louis, MO)). Ethical approval was obtained from the University of Otago Human Ethics Committee (Health): H14/046.

For MACS isolation of Vα7.2^+^ cells, PBMCs were stained with anti-Vα7.2 PE (BioLegend, San Diego, CA) and Vα7.2^+^ cells were isolated using anti-PE microbeads and MS columns (Miltenyi Biotec, Bergisch Gladbach, Germany) following manufacturer’s instructions.

### Bacteria and MAIT cell ligand

*E. coli* HB101 was grown overnight in Luria-Bertani broth (Sigma-Aldrich), fixed with 2% paraformaldehyde (PFA), washed with 1X PBS, and final bacterial suspension was counted using 123count eBeads (eBioscience, San Diego, CA). MR1 precursor ligand 5-amino-6-d-ribitylaminouracil (5-A-RU) was synthesised as previously described(45) and stored as single-use aliquots at -80°C. For each experiment, 5-A-RU and methylglyoxal (MG) (Sigma-Aldrich) were mixed at a molar ratio of 1:50 to form 5-A-RU/MG (referred to as 5-OP-RU), then diluted in milli-Q water to give desired concentrations.

### MAIT cell activation assays

PBMCs were seeded into 96-well U-bottom plates (Nunclonä Delta Surface, ThermoFisher, Waltham, MA) at 1x10^6^ cells/well unless otherwise stated.

PBMCs were treated with combinations of 5-OP-RU (1 nM, unless otherwise stated), 10 ng/mL IFN-α (Sigma-Aldrich), 1 ng/mL IFN-β (R&D Systems, Minneapolis, MN), 50 ng/mL IL-2, 50 ng/mL IL-7, 50 ng/mL IL-15, 10 ng/mL IL-21 (all from Peprotech, Rocky Hill, NJ), 10 ng/mL IL-12 (Miltenyi Biotec), 10 ng/mL IL-18 (R&D Systems), 10 bacteria per cell (BPC) fixed *E. coli* HB101, or a combination of 50 ng/mL PMA and 1 μg/mL ionomycin (Sigma-Aldrich). For blocking experiments 2.5 μg/mL anti-human/mouse/rat MR1 (Ultra-LEAF™ purified, 26.1, BioLegend), 1 μg/mL B18R (Invitrogen, Carslbad, CA), or 10 μg/mL anti-IL-21R (clone ATR-107, Absolute Antibody Ltd, Oxford, UK) were used.

For assessing CD107a expression, 1 mL anti-CD107a PE (H4A3, BioLegend) was added during treatment.

Samples were incubated for 6, 12, 24, 48, or 72 hrs at 37°C with 5% CO_2_. For assessing intracellular cytokines and cytotoxic molecules, 5 μg/mL brefeldin A (BioLegend) was added for the final 4 hrs.

### LEGENDplex^TM^ assay

MACS-isolated Vα7.2^+^ cells were seeded into 96-well U-bottom plates at 100,000 cells/well and either left untreated or stimulated with Dynabeads Human T-activator CD3/CD28 (ThermoFisher) (bead to cell ratio of 1:1) and 10 ng/mL IL-21 for 6 hrs. Experiment was performed in duplicate. Supernatants were harvested and stored at -80°C before analysis with either LEGENDplex™ HU Essential Immune Response Panel (13-plex) or LEGENDplex™ Human CD8/NK Panel (13-plex) (BioLegend) following manufacturer’s instructions. Samples were analysed using 4-laser Cytek™ Aurora and LEGENDplex™ Data Analysis Software Suite. Concentrations below the detection threshold were given the value of the limit of detection (denoted by a dotted line on each graph).

### Cytotoxic assays

The cytotoxic assay used in the current study is an adaptation of the FATAL assay(46) as described by Kurioka et al. 2015.(19) All assays were conducted in 96-well U-bottom plates.

BCL(47) (EBV-transformed B-cell lymphoblastoid cell line, kindly gifted by Paul Klenerman, University of Oxford) were stained with 1 μM carboxyfluorescein succinimidyl ester (CFSE) (eBioscience) or 1 μM Cell Trace Violet™ (CTV) (Invitrogen) following manufacturer’s instructions.

CFSE-stained BCL were treated with 10 nM 5-OP-RU for 1 hr or left untreated, then washed with R10 before addition of CTV-stained BCL (untreated) at a 1:1 ratio.

PBMCs or MACS-isolated Vα7.2^+^ cells pre-activated with IL-21 or vehicle (PBS) for 24 hrs were used as effector cells.

Treated Vα7.2^+^ cells were washed in R10 then added to mixed BCLs (CFSE- and CTV-stained) at a 2:1 ratio of Vα7.2^+^ to CFSE^+^ target BCL to give an approximate ratio of one MAIT cell to one CFSE^+^ BCL (given %MAIT of Vα7.2^+^ ranges from ∼50-85%). For PBMCs, the percentage of MAIT cells of live cells was first determined then used to calculate the number of PBMCs to add. For example, where % MAIT cells was 1% of live cells, 1x10^6^ PBMCs were added to 10,000 CFSE-stained BCL to maintain approximate ratio of 1 MAIT cell to 1 CFSE^+^ BCL.

Cells were incubated for 24 hrs; no Vα7.2^+^ cells or PBMCs were added to the no effector cell control (BCL only). For blocking MR1-TCR interaction, 2.5 μg/mL anti-MR1 (26.5) was added. After incubation, cells were washed, fixed in 2% PFA, stained, and assessed by flow cytometry.

The expected ratio (ER) of effector to target cells and % specific killing were calculated with the method below.(19) If the % specific killing was calculated to be negative, it was given an arbitrary value of 0.

Expected ratio (ER) = %CFSE^+^ cells / %CTV^+^ cells *(in BCL only control)* % Specific killing = 100 x [(ER x %CTV^+^ cells) - %CFSE^+^ cells]/(ER x %CTV^+^ cells)

### Flow cytometry

Cells were stained with Live/Dead near IR (Invitrogen) and antibodies against cell surface proteins for 25 min at 4°C. Cells were then either fixed/permeabilised with FoxP3 transcription factor staining buffer set (eBioscience) for 1 hr at room temperature (RT), or fixed with 2% PFA for 20 min at 4°C then treated with permeabilisation buffer (BioLegend). Cells were then stained with antibodies against TCR components (CD3, CD8, CD4, Vα7.2) and intracellular proteins for 25 min at RT. For flow cytometry analysis of B cells, cells were incubated with Human TruStain FcX™ (BioLegend) for 10 min at 4°C before surface staining. Cells were washed in FACS buffer (PBS + 2% FBS) for all wash steps.

For pSTAT staining, cells were surface stained, washed, resuspended in R10, then treated with cytokines for 15 min at 37°C. Cells were fixed immediately with 2% PFA for 20 min at 37°C, permeabilised with cold True Phos™ buffer (BioLegend) for 1 hr at -20°C, then stained for pSTAT proteins for 30 min at RT.

Antibodies used were: CD161 APC or Brilliant Violet™ (BV) 605 (HP-3G10, BioLegend), CD69 FITC or PerCP-Cy5.5 (FN50, BioLegend), CD178 (FasL) PE-Cy7 (NOK-1, BioLegend), CD360 (IL-21R) PE (17A12, BioLegend), CD19 FITC (HIB19, BioLegend), CD3 PE-Cy7 (UCHT1, BioLegend), CD3 BV510 (OKT3, BioLegend), CD8 eFluor450 (RPA-18, eBioscience), TCR Vα7.2 PE, PE-Cy7, Alexa-Fluor700 (AF700), or BV711 (3C10, BioLegend), CD4 AF700 (SKA3, BioLegend), CD4 FITC (RPA-T4, BioLegend), GrzB FITC or PE-Cy7 (QA16AO2, BioLegend), GrzA PE (CB9, BioLegend), perforin PerCP-Cy5.5 (B-D48, BioLegend), TNF-α FITC (Mab11, BioLegend), IFN-γ PerCP-Cy5.5 (4S.B, BioLegend), IL-17A BV605 (BL168, BioLegend), granulysin PE-Cy7 (DH2, BioLegend), IL-21 PE (4BG1, BioLegend), Blimp1 PE-CF594 (6D3, BD Biosciences), T-bet PE-Cy7 (4B10, BioLegend), pSTAT1 (Tyr701) PE (A17012A, BioLegend), pSTAT3 (Tyr705) PE-Cy5 (13A3-1, BioLegend), pSTAT5 (Tyr694) APC (A17016B, BioLegend).

Samples were acquired on a 3-laser FACSCanto™ II or 4-laser BD LSRFortessa™ (BD Biosciences), or 4-laser Cytek™ Aurora (Cytek Biosciences, Fremont, CA). Data were analysed on FlowJo™ V10.10.0 (TreeStar, Ashland, OR). MAIT cells were defined by expression of TCR Vα7.2 and high expression of CD161.(5) Gating strategies can be found in Supporting Information Fig. 1-5.

**Figure 1:**
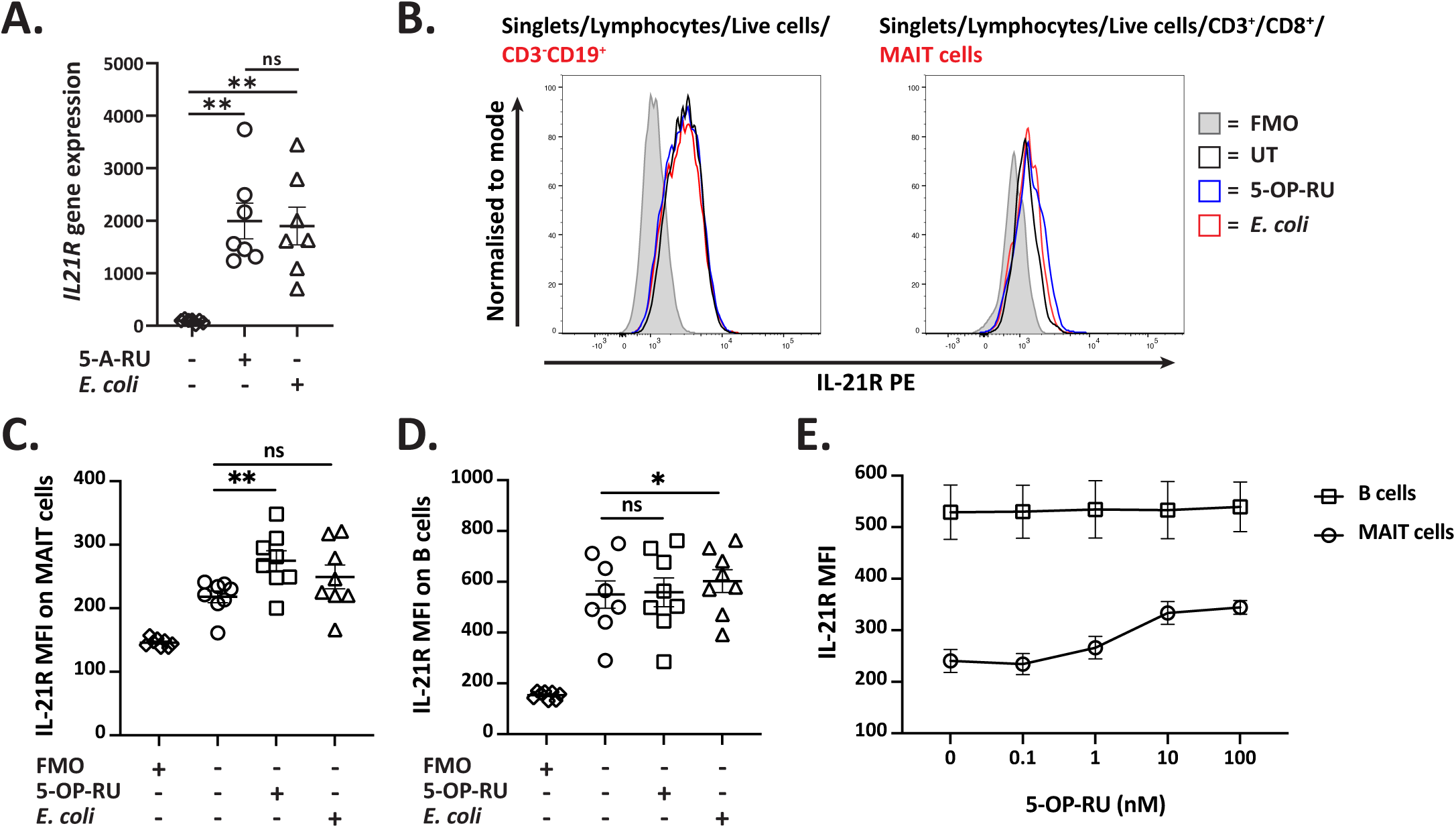
Activated MAIT cells upregulate IL-21R expression. (A) *IL21R* gene counts from RNA-seq of MAIT cells stimulated with 5-A-RU or *E. coli* for 6 hrs. (B-D) PBMCs were treated with 5-OP-RU or *E. coli* (B-D) or different concentrations of 5-OP-RU (E) for 6 hrs, then IL-21R MFI on MAIT (CD3**^+^**CD8**^+^**CD161**^++^**Vα7.2**^+^**) and B cells (CD3**^-^**CD19⁺) was assessed by flow cytometry; 0 nM represents untreated (E). (B) Representative histograms showing IL-21R expression on B and MAIT cells. Each biological replicate and the mean ± SEM (n=7-8) (A, C-D) or the mean ± SEM (n=8) (E) from two independent experiments are shown. (A,C-D) Data were analysed with repeated measures ANOVA with Sidak’s multiple comparison post-test. MFI = median fluorescence intensity. ns = no significance, *p<0.05, **p<0.01.

### Bioinformatic analysis

For analysis of gene expression of *IL21* and *IL21R* by MAIT cells, gene count data was obtained from our previous RNA-seq study.(45)

### Statistical analysis

DCata were analysed using GraphPad Prism version 9 (GraphPad Software, San Diego, CA). First, data were tested for normality with Shapiro-Wilk test. For normally distributed data, multiple groups were compared with repeated measures one-way or two-way analysis of variance (ANOVA) with Sidak’s multiple comparison test. For data not normally distributed, multiple groups were compared with Friedman’s test with Dunn’s multiple comparison test. p<0.05 was considered significant.

## Results

### Activated MAIT cells express IL-21R

Previous research has shown that IL-21 enhances cytotoxic response of neutrophils, NK cells, and CD8^+^ T cells(42-44) and that MAIT cells express IL-21R.(41) Therefore, we hypothesised that IL-21 can modulate MAIT cell TCR-mediated cytotoxic potential.

To confirm expression of IL-21R by MAIT cells, we revisited our previous RNAseq dataset (45) and assessed *IL21R* gene expression (Fig. 1A). Expression of *IL21R* was significantly higher in MAIT cells stimulated with *E. coli* and pre-cursor TCR ligand 5-A-RU (p<0.01) compared to resting MAIT cells, with 5-A-RU inducing the highest expression (Fig. 1A).

To confirm surface expression, PBMCs were stimulated with 5-OP-RU or *E. coli* and expression of IL-21R was assessed by flow cytometry (Fig. 1B-D). Given that IL-21R is highly expressed on B cells,(48) they were included as a positive control. MAIT cells stimulated with *E. coli* or 5-OP-RU upregulated IL-21R expression, as determined by median fluorescence intensity (MFI), although this was only statistically significant for 5-OP-RU (p<0.01) (Fig. 1C). As expected, IL-21R was expressed on B cells and MFI was not impacted by 5-OP-RU (Fig. 1D). To further confirm the role of TCR stimulation on IL-21R expression by MAIT cells, PBMCs were treated with different concentrations of 5-OP-RU (Fig. 1E). IL-21R MFI on MAIT cells, but not B cells, increased with increasing concentrations of 5-OP-RU (Fig. 1E), suggesting a dose-dependent response and the potential to respond to IL-21.

### IL-21 enhances TCR-dependent MAIT cell cytotoxic potential

To evaluate the effect of IL-21 on TCR-activated MAIT cells, PBMCs were stimulated with combinations of IL-21, 5-OP-RU, and anti-MR1 (αMR1) for 6 hrs (Fig. 2A-F). IL-21 treatment alone significantly enhanced perforin expression on MAIT cells (p<0.0001, Fig. 2D) but had no or only minor effects on other markers (Fig. 2A-C and 2E-F). In 5-OP-RU-activated MAIT cells, IL-21 further enhanced expression of CD69, CD107a, and GrzB in an MR1-dependent manner (Fig. 2A-B and 2E) but did not impact 5-OP-RU-induced downregulation of GrzA (Fig. 2F). No biologically significant expression of FasL or granulysin was observed on MAIT cells, despite modest increases in FasL MFI (Fig. 2C and Supporting Information Fig. 6A). Additionally, IL-21 had negligible effects on IFN-γ, TNF-α, and IL-17A expression by 5-OP-RU-stimulated MAIT cells (Supporting Information Fig. 6A and 6B-C).

**Figure 2.**
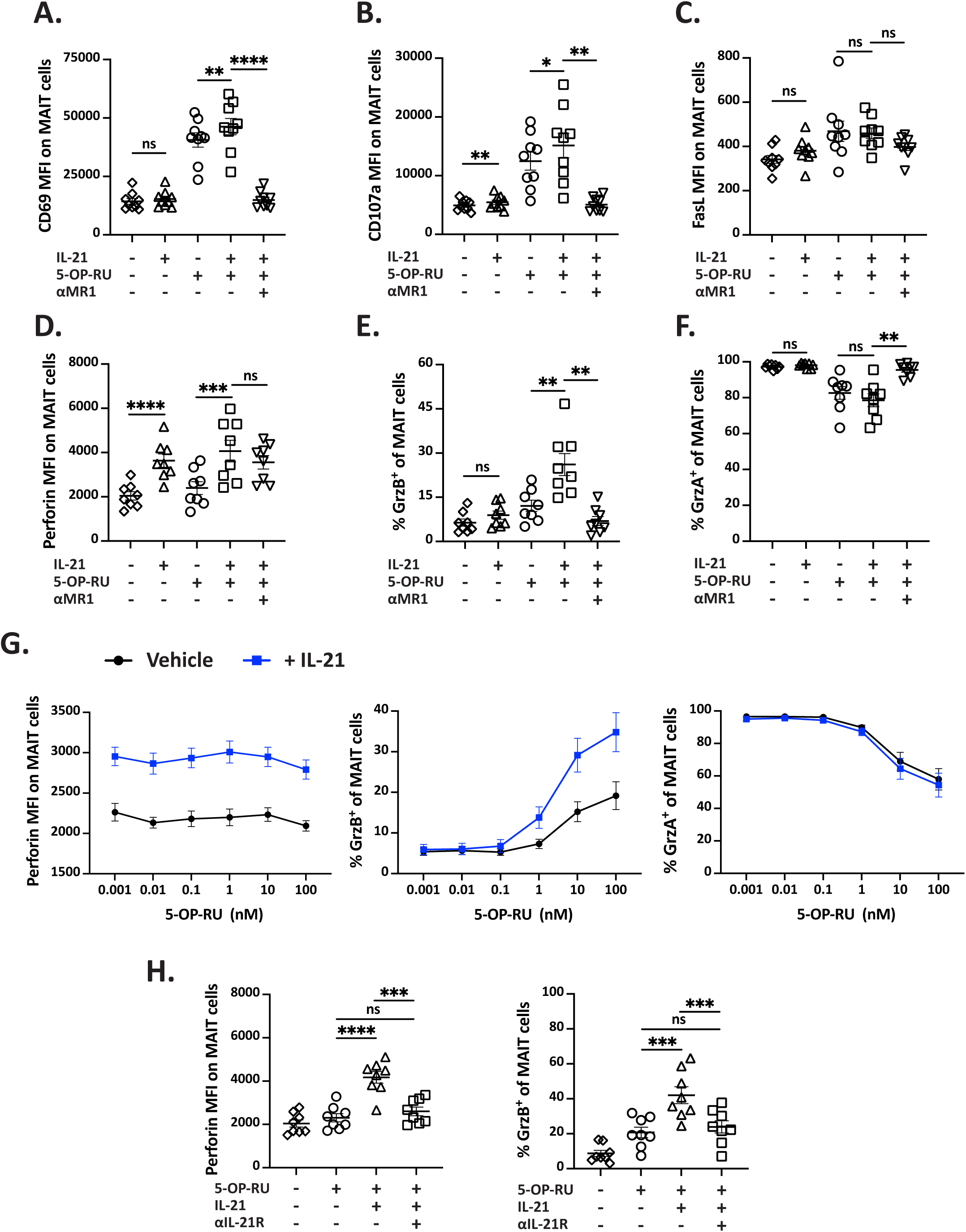
IL-21 enhances TCR-dependent MAIT cell cytotoxic potential. (A-F) PBMCs were treated with combinations of IL-21, 5-OP-RU, and αMR1 for 6 hrs. MFI of CD69 (A) CD107a (B), FasL (C), perforin (D) and the frequency of MAIT cells expressing GrzB (E) and GrzA (F) were assessed by flow cytometry. (G) PBMCs were treated with different concentrations of 5-OP-RU with PBS (vehicle) or IL-21 for 6 hrs then assessed for expression of perforin, GrzB and GrzA. (H) PBMCs were treated with combinations of IL-21, 5-OP-RU, and αIL-21R for 6 hrs and assessed for expression of perforin and GrzB. For assessing cytotoxic molecules, (D-H) brefeldin A was added for the final 4 hrs. Each biological replicate (n=9 (A-C), n=8 (D-F, H) and the mean ± SEM is shown. Alternatively, data is shown as mean ± SEM (n=9) (G). All data was obtained from two or three independent experiments. Data were analysed with repeated measures one-way ANOVA with Sidak’s multiple comparison post-test or Friedman’s test with Dunn’s multiple comparison post-test. MFI = median fluorescence intensity. ns = no significance, *p<0.05,**p<0.01,***p<0.001, ****p<0.0001.

To further illustrate the combined effect of IL-21 and 5-OP-RU, PBMCs were stimulated with IL-21 and varying concentrations of 5-OP-RU (Fig. 2G). IL-21 enhanced perforin expression independently of 5-OP-RU concentration, while its effect on GrzB was only evident at 5-OP-RU concentrations of 1 nM or more. In contrast, IL-21 did not impact GrzA expression (Fig. 2G). Furthermore, IL-21-mediated enhancement of perforin and GrzB could be completely inhibited by addition of anti-IL-21R (αIL-21R) (Fig. 2H).

To determine whether IL-21 acts directly on MAIT cells, MACS-isolated Vα7.2^+^ cells (>95% purity, Supporting Information Fig. 3) were stimulated with IL-21 and/or anti-CD3/CD28 (αCD3/CD28) for 6 hrs (Fig. 3). Cell culture supernatants were analysed by LEGENDplexä (Fig. 3C-N) and the cells assessed for GrzB expression by flow cytometry (Fig. 3A-B). IL-21 + αCD3/CD28 significantly increased the percentage of MAIT cells expressing GrzB compared to stimulation with αCD3/CD28 alone (p<0.01, Fig. 3A) but only a small, non-significant increase in GrzB concentration was detected in the culture supernatant (Fig. 3C). IL-21 did not impact production of GrzA, perforin, soluble FasL (sFasL) or granulysin (Fig. 3D-G). Interestingly, IL-21 downregulated IL-2 (p<0.05) and TNF-α (p<0.01) production by αCD3/CD28-stimulated Vα7.2^+^ cells, but had no effect on IFN-γ and IL-17A (Fig. 3H-K). IL-8, IL-10, and CXCL10 were also detected in culture supernatants but no effect of IL-21 on their expression was observed (Fig. 3L-N).

**Figure 3.**
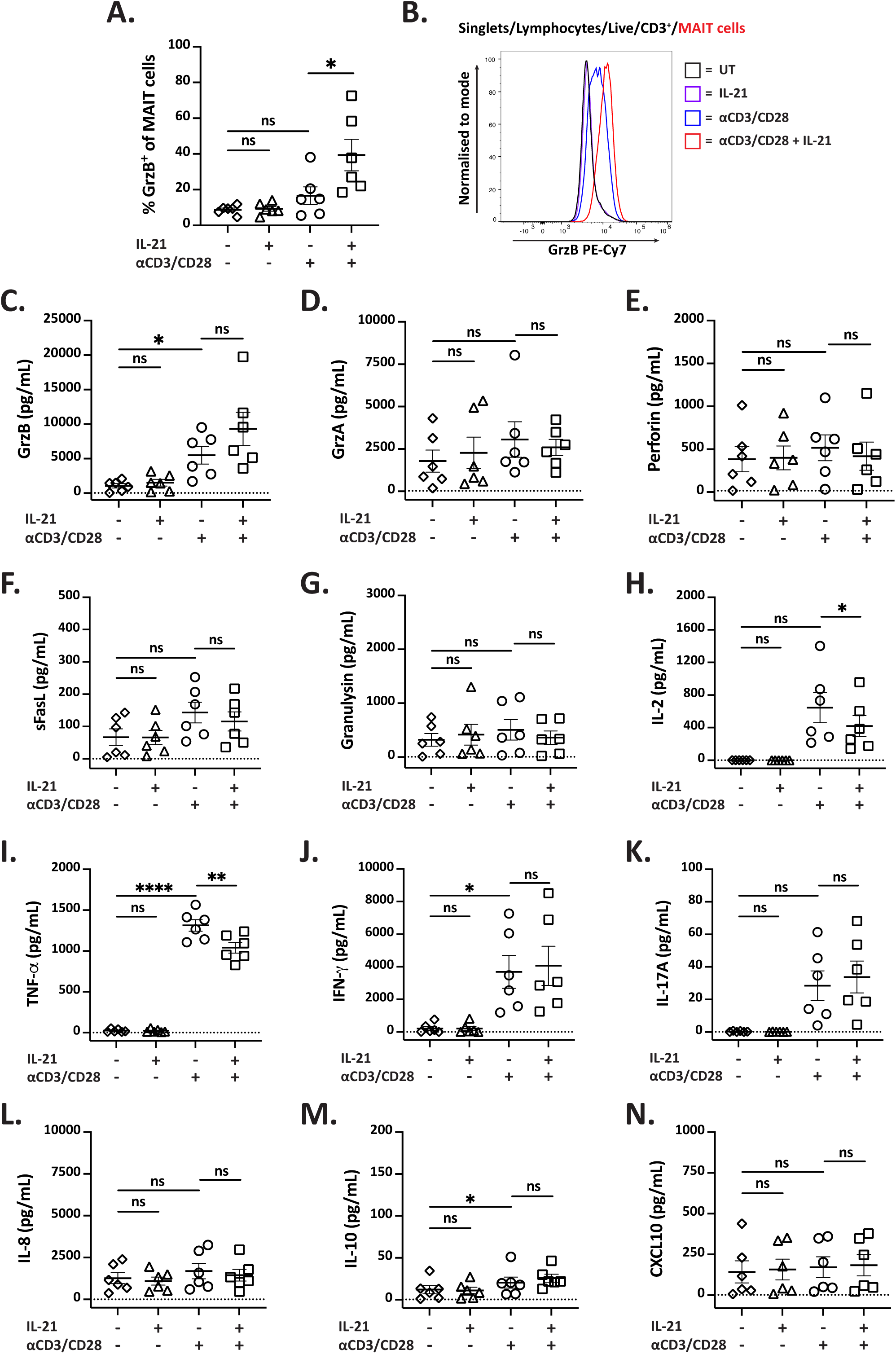
IL-21 acts directly on MAIT cells to enhance GrzB production but has limited impact on cytokine production. (A-N) MACS-isolated Vα7.2^+^ cells were treated with IL-21 and αCD3/CD28 beads for 6 hrs. Culture supernatants were analysed using LEGENDplex™ (C-N) and remaining cells assessed for GrzB expression by flow cytometry (A). Representative histogram showing GrzB on MAIT cells is also shown (B). Concentrations of GrzB (C), GrzA (D), perforin (E), sFasL (F), granulysin (G), IL-2 (H), TNF-α (I), IFN-γ (J), IL-17A (K), IL-8 (L), IL-10 (M), and CXCL10 (N) were obtained using LEGENDplex™. Each biological replicate (n=6) and the mean ± SEM from two independent experiments are shown. Data were analysed with repeated measures one-way ANOVA with Sidak’s multiple comparison post-test or Friedman’s test with Dunn’s multiple comparison post-test. MFI = median fluorescence intensity. ns = no significance,* p<0.05,** p<0.01,**** p<0.0001.

Overall, IL-21 enhances GrzB expression by TCR-activated MAIT cells but has minimal impact on cytokine production.

### IL-21 enhances TCR-independent MAIT cell cytotoxic potential

To assess whether MAIT cell responses to IL-21 are time-dependent, PBMCs were treated with IL-21 for varying durations over a 72-hr period (Supporting Information Fig. 7A-D). Perforin expression peaked after 6 hrs then declined over time, while GrzB expression increased over 48 hrs (Supporting Information Fig. 7A-B). Both GrzA and granulysin expression increased over the 72-hr period (Supporting Information Fig. 7C-D). These findings suggest that IL-21 differentially regulates expression of cytotoxic molecules by MAIT cells over time.

Given the time-dependent effects of IL-21 on MAIT cells, we investigated whether IL-21 could enhance TCR-independent MAIT cell activation by IL-12 and IL-18, which occurs later than TCR-dependent activation.(26) PBMCs were treated with IL-21 and a combination of IL-12 and IL-18 for 24 hrs (Fig. 4A-F). IL-21 had no impact on CD69, CD107a, FasL or GrzA expression without IL-12/IL-18 co-stimulation (Fig. 4A-C, F). In contrast to observations at 6 hrs (Fig. 2), IL-21 significantly enhanced GrzB expression on MAIT cells (p<0.001, Fig. 4E), but its effect on perforin was modest (Fig. 4D). When combined with IL-12/IL-18, IL-21 further upregulated expression of all cytotoxic markers, but had no additional effect on CD69 MFI (Fig. 4A-F). Thus, IL-21 enhances expression of cytotoxic molecules by MAIT cells in combination with IL-12 and IL-18.

**Figure 4.**
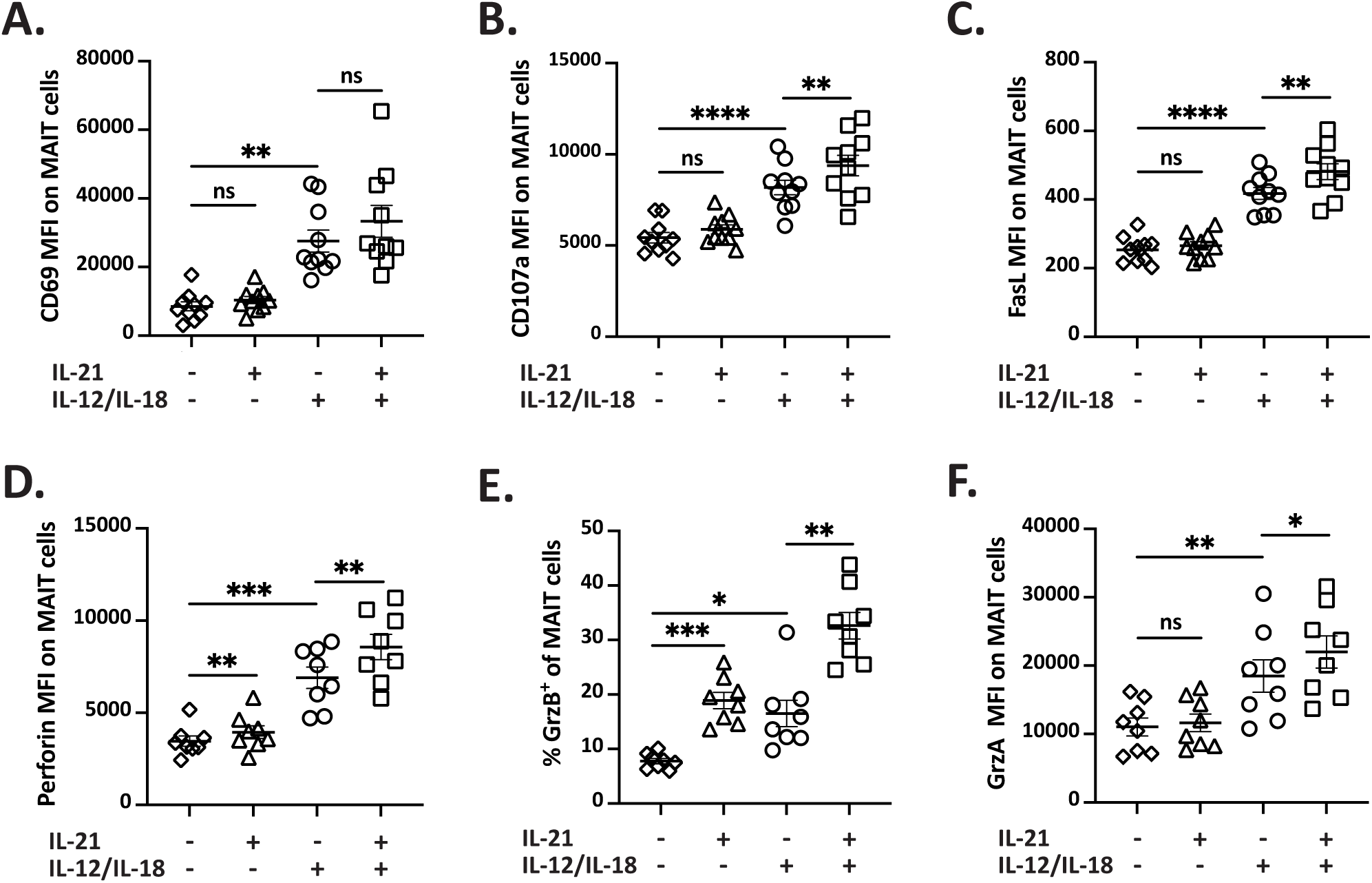
IL-21 enhances TCR-independent MAIT cell cytotoxic potential. (A-F) PBMCs were treated with IL-21 and/or IL-12 and IL-18 for 24 hrs then MFI of CD69 (A), CD107a (B), FasL (C), perforin (D), and GrzA (F) and the frequency of MAIT cells expressing GrzB (E) were assessed by flow cytometry; (D-F) brefeldin A was added for the final 4 hrs. Each biological replicate (n=10 (A-C), n=8 (D-F)) and the mean ± SEM from two or three independent experiments are shown. Data were analysed with repeated measures one-way ANOVA with Sidak’s multiple comparison post-test or Friedman’s test with Dunn’s multiple comparison post-test. MFI = median fluorescence intensity. ns = no significance, *p<0.05,**p<0.01,***p<0.001,****p<0.0001.

### Additive effects of IL-21 and other cytokines on TCR response of MAIT cells

In addition to triggering TCR-independent activation, cytokines, such as T1-IFNs, enhance TCR-dependent MAIT cell activation.(29) To assess whether IL-21 synergises with T1-IFNs, PBMCs were treated with 5-OP-RU, IL-21, and sub-optimal concentrations of IFN-α or IFN-β for 6 hrs (Supporting Information Fig. 8A-D). The individual effects of IL-21 and T1-IFNs on MAIT cells were comparable, and when combined, only a modest increase in GrzB and perforin expression was observed (Supporting Information Fig. 8A-D), suggesting they act independently but may have an additive effect on MAIT cells. A similar additive effect was observed when PBMCs were stimulated with 5-OP-RU and IL-21 in combination with IL-2, IL-7, or IL-15 for 6 hrs (Supporting Information Fig. 8G-H). Treatment with any cytokine modestly enhanced GrzB expression by MAIT cells, but when combined with IL-21, the frequency of MAIT cells expressing GrzB significantly increased (Supporting Information Fig. 8H). Additionally, in the absence of 5-OP-RU, perforin expression, but not GrzB, was significantly enhanced by IL-21, while other cytokines had minimal effect (Supporting Information Fig. 8E-F). Overall, at 6 hrs, IL-21 significantly upregulates GrzB and perforin expression by MAIT cells, and this response can be further boosted by the presence of T1-IFNs or cytokines IL-2, IL-7, and IL-15.

### Expression of Blimp-1 and pSTAT3 are upregulated by IL-21

IL-21 has a response element downstream of the *prdm1* (Blimp-1) gene(49) and induces expression of transcription factors Blimp-1 and T-bet.(50, 51) Given Blimp-1 and T-bet are associated with enhanced effector response of MAIT cells,(19) we hypothesised that MAIT cell response to IL-21 may involve increased expression of Blimp-1 and T-bet.

To evaluate this, PBMCs were stimulated with IL-21, with or without 5-OP-RU, and Blimp-1 and T-bet expression on MAIT cells was assessed by flow cytometry (Fig. 5A-B). Neither IL-21 nor 5-OP-RU had an independent effect, but when combined, they significantly enhanced Blimp-1 MFI compared to unstimulated MAIT cells (p<0.0001, Fig. 5A). Interestingly, treatment with IL-21 alone resulted in a modest decrease in T-bet MFI, while 5-OP-RU significantly upregulated T-bet (Fig. 5B). Therefore, Blimp-1 signalling may be involved in MAIT cell responses to 5-OP-RU and IL-21.

**Figure 5:**
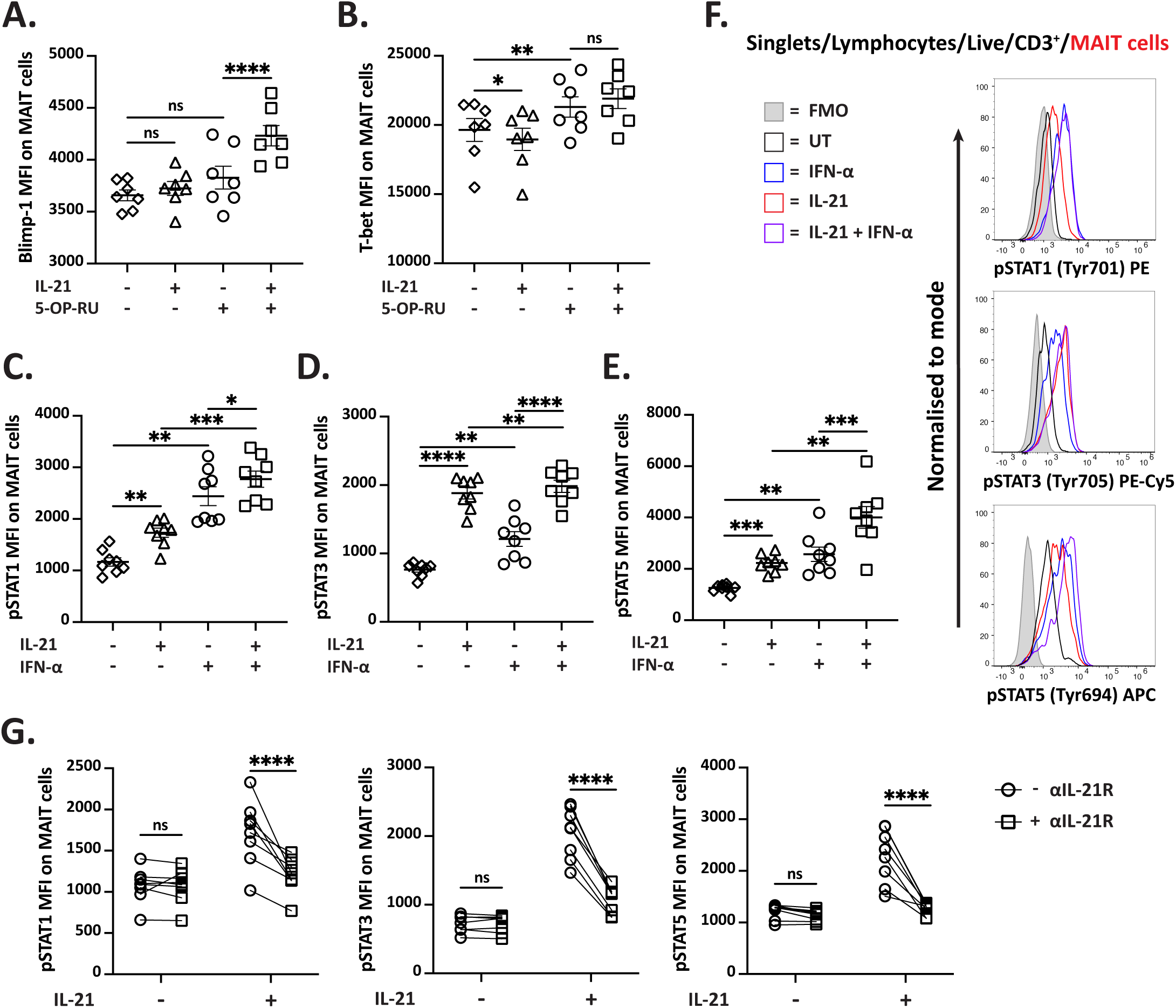
IL-21 activates multiple STAT proteins in MAIT cells. (A-B) PBMCs were treated with IL-21 and/or 5-OP-RU for 6 hrs then MFI of Blimp-1 (A) and T-bet (B) on MAIT cells was assessed by flow cytometry. (C-F) PBMCs were treated with IL-21 and/or IFN-α for 15 min then MFI of pSTAT1 (C), pSTAT3 (D), and pSTAT5 (E) in MAIT cells was assessed; representative histograms of pSTAT expression are shown (F). (G) PBMCs were treated with IL-21 with or without αIL-21R for 15 min then MFI of pSTAT1, pSTAT3, and pSTAT5 was assessed. Each biological replicate (n=7 (A-B), n=8 (C-G)), and the mean ± SEM from two independent experiments are shown. Data were analysed with either repeated measures one-way or two-way ANOVA with Sidak’s multiple comparison post-test. MFI = median fluorescence intensity. ns = no significance,*p<0.05,** p<0.01,***p<0.001,****p<0.0001.

IL-21 induces phosphorylation of STAT1, STAT3, and STAT5 in T cells(38-40, 52) and STAT3 signalling is important for maintenance of MAIT cell numbers in humans.(41) To further elucidate the mechanisms underlying MAIT cell responses to IL-21, PBMCs were stimulated with IL-21 and assessed for expression of pSTAT proteins in MAIT cells (Fig. 5C-F); IFN-a was included for comparison. Representative flow plots for each treatment are also shown (Fig. 5F). Both IFN-a and IL-21 significantly induced phosphorylation of STAT1, STAT3, and STAT5 in MAIT cells (Fig. 5C-E), with pSTAT3 exhibiting the strongest upregulation by IL-21 (p<0.0001, Fig. 5D). Expression of all pSTAT proteins further increased when IL-21 and IFN-a were combined, compared to treatment with either cytokine alone (Fig. 5C-E), highlighting the additive effect of T1-IFNs and IL-21 on MAIT cells. Additionally, phosphorylation of STAT proteins was inhibited when PBMCs were pre-treated with αIL-21R before stimulation with IL-21 (Fig. 5G), confirming this response is specific to IL-21.

In summary, IL-21 stimulates multiple STAT pathways in MAIT cells, but how this correlates with MAIT cell response to bacteria remains to be determined.

### GrzB expression by *E. coli*-stimulated MAIT cells involves IL-21 signalling

We have shown that adding exogenous IL-21 modulates MAIT cell cytotoxic potential, but it is unclear whether IL-21 is important for MAIT cell responses to a physiological stimulus, such as bacteria. To assess this, we treated PBMCs with *E. coli* and αIL-21R (Fig. 6A-D). Addition of αIL-21R resulted in a significant decrease in GrzB expression by *E. coli-*treated MAIT cells (p<0.0001, Fig. 6A). The effect of blocking IL-21R on perforin expression was less clear, however, a decrease in perforin MFI was observed with addition of αIL-21R (p<0.05, Fig. 6B). As expected, αIL-21R did not impact IFN-γ or TNF-α expression by MAIT cells (Fig. 6C-D).

**Figure 6:**
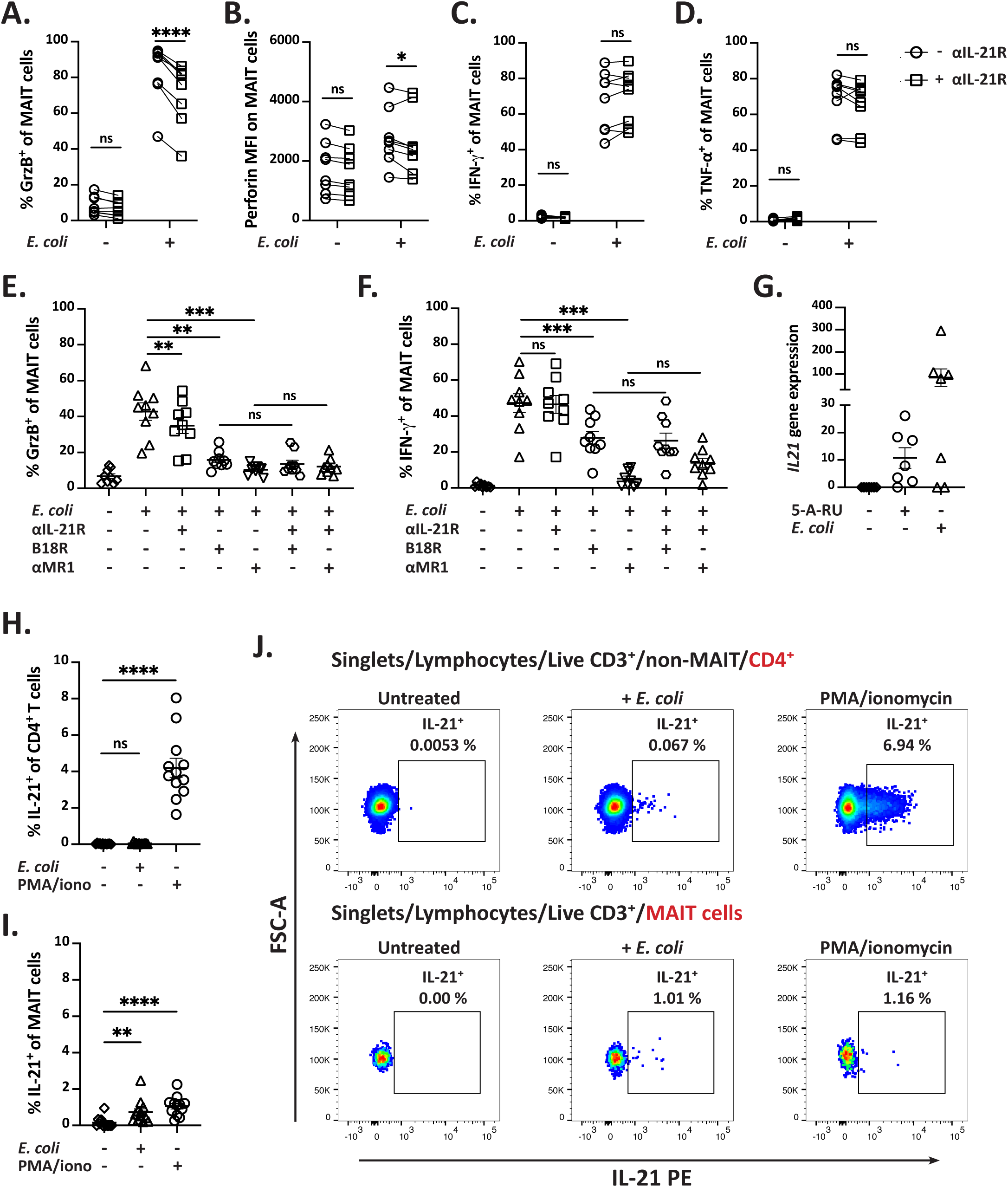
GrzB-expression by *E. coli*-stimulated MAIT cells involves IL-21 signalling. (A-D) PBMCs were treated with *E. coli* with or without αIL-21R for 6 hrs. Perforin MFI (B) and the frequency of MAIT cells expressing GrzB (A), IFN-γ (C), and TNF-α (D) were assessed by flow cytometry. (E-F) PBMCs were treated with *E. coli* and combinations of αIL-21R, B18R, and αMR1 for 6 hrs then frequency of MAIT cells expressing GrzB (E) and IFN-γ (F) was assessed. (G) *IL21* gene counts from RNA-seq of MAIT cells stimulated with 5-A-RU or *E. coli* for 6 hrs. (H-J) PBMCs were treated with *E. coli* or PMA/ionomycin for 6 hrs and %IL-21^+^ of CD4⁺ T cells (H) and MAIT cells (I) was assessed. (J) Representative flow plots for each treatment are shown; gates defining IL-21^+^ were set using untreated samples. (A-F, H-J) Brefeldin A was for the final 4 hrs. Each biological replicate (n=8 (A-D), n=9 (E-F), n=7 (G), n=13 (H-I)) and the mean ± SEM from two or three independent experiments are shown. Data were analysed with repeated measures one-way or two-way ANOVA with Sidak’s multiple comparison post-test or Friedman’s test with Dunn’s multiple comparison post-test. MFI = median fluorescence intensity. ns = no significance, *p<0.05, **p<0.01, ***p<0.001, ****p<0.0001.

To compare the effects of blocking IL-21-, TCR-, and T1-IFN-signalling during MAIT cell activation with *E. coli*, PBMCs were treated with *E. coli*, and combinations of αIL-21R, αMR1, and B18R, a vaccinia virus protein that neutralises T1-IFNs(53) (Fig. 6E-F), and GrzB and IFN-γ expression was assessed. Only GrzB expression decreased with αIL-21R (p<0.01, Fig. 6E), whereas B18R treatment significantly reduced both GrzB and IFN-γ expression (p<0.01 and p<0.001, respectively, Fig. 6E-F). As expected, the strongest inhibition of GrzB and IFN-γ expression by MAIT cells was observed with αMR1 (both p<0.001, Fig. 6E-F). Combining αIL-21R with either B18R or αMR1 provided no additional blocking effect for either GrzB or IFN-γ, reiterating that TCR and T1-IFNs are the dominant signals driving early MAIT cell response to *E. coli*.

These findings suggest that IL-21 was present in our PBMC culture and contributed to MAIT cell cytotoxic responses to *E. coli.* CD4^+^ T cells and iNKT cells are dominant producers of IL-21,(32, 33) and MAIT cells also produce IL-21 in some contexts.(34-37) However, it remains unclear whether IL-21 was produced by our PBMC culture. First, we revisited our previous RNAseq dataset(45) to analyse *IL21* gene counts on MAIT cells stimulated with 5-A-RU or *E. coli* (Fig. 6G). PBMC MAIT cells expressed minimal *IL21*, supporting findings from previous studies,(34, 37) although a modest increase in *IL21* expression was observed with *E. coli* stimulation (Fig. 6G).

To demonstrate production of IL-21 by flow cytometry, PBMCs were treated with *E. coli* or PMA/ionomycin (as a control) for 6 hrs and IL-21 expression on CD4^+^ T cells and MAIT cells was assessed (Fig. 6H-J). Limited production of IL-21 by *E. coli*-stimulated CD4^+^ T cells and MAIT cells was observed, as only a few cells exhibiting positive staining (Fig. 6H-J). While the IL-21^+^ MAIT cells increased, percentage was low (Fig. 6I). Overall, CD4^+^ T cells and MAIT cells in PBMCs produce little IL-21 in response to 6 hr-stimulation with *E. coli*.

### IL-21 enhances direct killing by MAIT cells

We have demonstrated that IL-21 enhances TCR-dependent and TCR-independent production of cytotoxic molecules by MAIT cells, however, whether IL-21 enhances MAIT cell killing remains to be determined.

We adapted a previously published cytotoxic assay method(19) using PBMCs as effector cells to assess MAIT cell killing of CFSE-stained BCLs (B-cell lymphoblastoid cell line) treated with 5-OP-RU (Fig. 7A). PBMCs were pre-treated with PBS (vehicle) or IL-21 for 24 hrs before addition to untreated or 5-OP-RU-treated BCLs (Fig. 7C-D); representative flow plots showing %CFSE^+^ vs CTV^+^ BCLs for each treatment are also shown (Fig. 7B). IL-21 significantly enhanced killing of 5-OP-RU-treated BCLs compared to untreated PBMCs (p<0.05, Fig. 7C-D). Interestingly, IL-21 only enhanced GrzB expression on MAIT cells incubated with untreated BCLs (Fig. 7D). Mean percentage specific killing by MAIT cells was between 30-40% (Fig. 7C), consistent with previous findings.(19)

**Figure 7:**
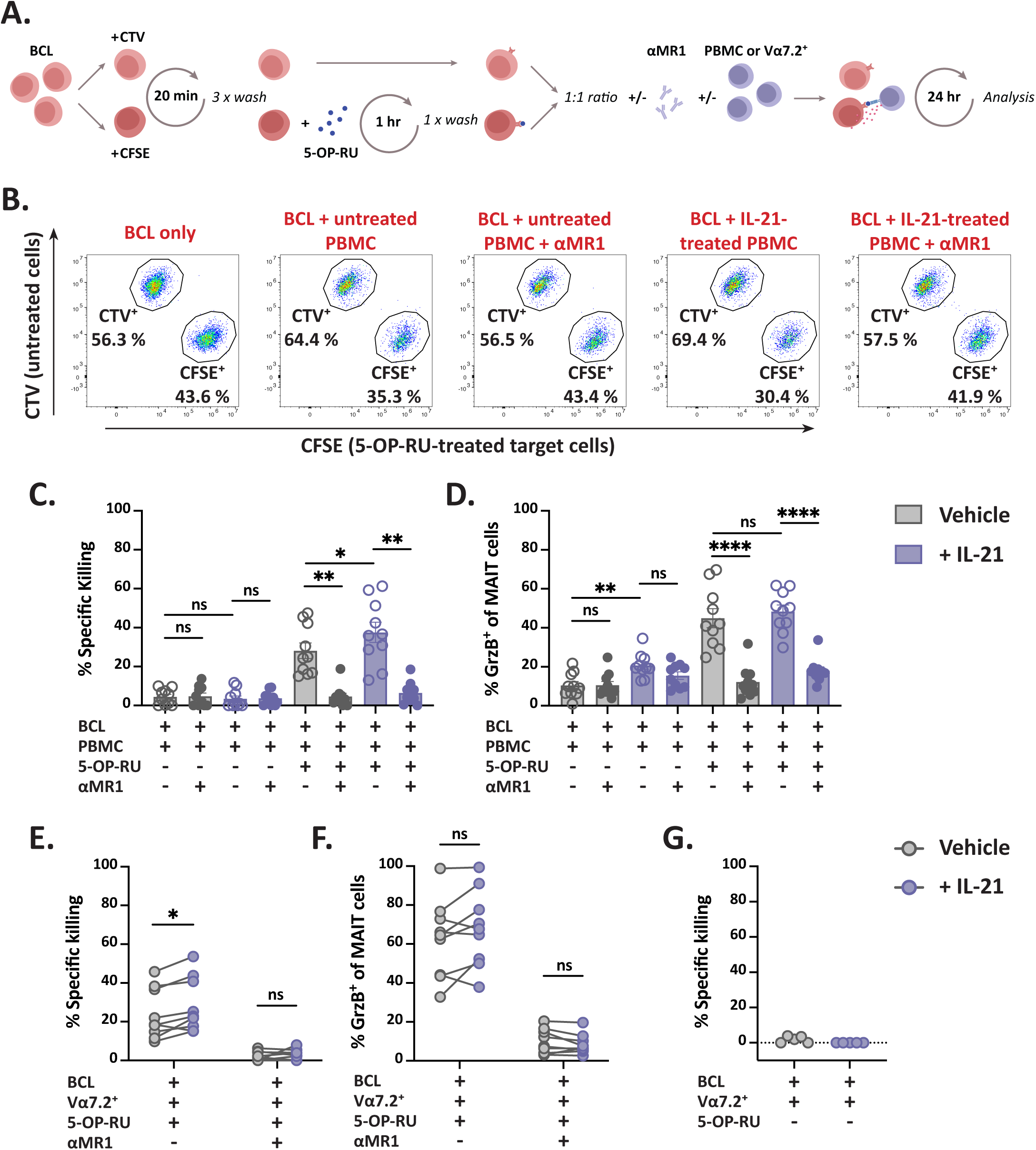
IL-21 augments MAIT cell killing ability. (A) Diagram of assay for assessing MAIT cell killing using PBMC or Vα7.2^+^ cells as effector cells. CFSE-stained BCLs were treated with 5-OP-RU or left untreated for 1 hr then washed before addition of untreated CTV-stained BCLs. PBMCs or Vα7.2^+^ were treated with PBS (vehicle) or IL-21 for 24 hrs before addition to BCLs with or without αMR1; cells were incubated for 24 hrs. (B) Representative flow plots showing frequencies of CFSE^+^ and CTV^+^ BCLs for each treatment. Percentage specific killing (C) and the frequency of MAIT cells expressing GrzB (D) were assessed by flow cytometry. (E-G) Vα7.2^+^ were added to 5-OP-RU-(E-F) or untreated-BCL (G) with or without αMR1 and percentage specific killing (E, G) and the frequency of MAIT cells expressing GrzB (F) were assessed. Each biological replicate (n=10 (C-D), n=9 (E-F), n=5 (G)) from two or four independent experiments are shown. Data were analysed with two-way ANOVA with Sidak’s multiple comparison post-test. MFI = median fluorescence intensity. ns = no significance, *p<0.05, **p<0.01, ****p<0.0001.

To further investigate the direct effect of IL-21 on MAIT cell cytotoxicity, the assay was repeated using Vα7.2^+^ cells as effector cells (Fig. 7E-G). IL-21 resulted in a small, but significant enhancement of specific killing by Vα7.2^+^ cells (p<0.05, Fig. 7E), reflecting the results seen with PBMCs. Killing was abrogated by αMR1, and IL-21 conferred no ability to kill untreated BCLs (Fig. 7G), reiterating that MAIT cell killing requires TCR-ligand interaction. There was no difference in GrzB expression between MAIT cells pre-treated with PBS or IL-21 (Fig. 7F). In conclusion, priming MAIT cells with IL-21 augments their ability to kill 5-OP-RU-treated target cells in a TCR-dependent manner.

## Discussion

IL-21 is a known inducer of cytotoxic responses. In CD8^+^ T cells, perforin expression is induced by IL-21,(54) and IL-21 upregulates GrzB expression in multiple cell types, including neutrophils,(42) iNKT cells(33), CD8^+^ T cells,(44) dendritic cells (DC),(55) B cells,(56) and NK cells.(57) Given MAIT cells express IL-21R,(41) we hypothesised that IL-21 modulates MAIT cell cytotoxicity. Here, we demonstrate IL-21 augments the cytotoxic potential of activated MAIT cells. Using *in vitro* PBMC assays, IL-21 significantly enhanced expression of GrzB and perforin by TCR- and cytokine-activated MAIT cells, and improved their ability to kill 5-OP-RU-treated BCLs. Interestingly, while IL-21 upregulated cytotoxic molecule expression, it had minimal impact on expression of IFN-γ, TNF-α, or IL-17A, suggesting a specific role in priming MAIT cells for cytotoxic functions.

Cytotoxic molecule expression by MAIT cells is a time-dependent process and their cytotoxic granule content differs between early TCR-dependent and late TCR-independent activation.(45) In the present study, we primarily assessed MAIT cell responses at 6 hrs, a time-point where activation relies on TCR-MR1 interactions.(47) Within this time-frame, IL-21 enhanced expression of GrzB by TCR-activated MAIT cells, while perforin expression was upregulated by IL-21 independently of TCR stimulation. These findings are consistent with previous observations that GrzB expression in MAIT cells is triggered by both TCR-ligation(45) and cytokines,(22, 26, 27) whereas perforin expression is induced independently of TCR signalling.(19) At 6 hrs, IL-21 alone did not impact expression of GrzA, GrzB, or granulysin, but an increase in their MFI over 72 hr was observed, suggesting a time-dependent response to IL-21. Consistent with previous reports,(19, 45) we did not observe biologically significant FasL surface expression on unstimulated MAIT cells. However, we observed a modest increase in FasL MFI following 24 hr stimulation with IL-12/IL-18 and IL-21, which aligns with a previous study demonstrating 24 hr stimulation with IL-12/IL-18 increases FasL surface expression on MAIT cells.(45) Overall, these results suggest that extended exposure to IL-21 influences expression of cytotoxic molecules by MAIT cells.

Although MAIT cell killing is TCR-dependent,(17, 19) priming MAIT cells for extended periods with additional signals enhances their killing ability. Pre-treatment of MAIT cells with IL-7 significantly augments their ability to kill *E. coli*-infected cell lines.(22, 58) Similarly, Kurioka et al.(19) demonstrated that MAIT cell killing was enhanced by priming MAIT cells with *E. coli* over a 6 day period. Using a modified version of their cytotoxic assay, we showed that priming PBMC and Vα7.2^+^ cells with IL-21 for 24 hrs resulted in a small, but significant, enhancement in MAIT cell killing of 5-OP-RU-treated BCLs. It is possible that prolonged exposure to IL-21 may induce more substantial changes in MAIT cell cytotoxic granule content that could further enhance their killing ability, however, further work is needed to evaluate this.

Previous studies have demonstrated that IL-21 induces expression of Blimp-1 and T-bet,(49-51) and GrzB production is associated with Blimp-1 and T-bet expression in human MAIT cells.(19) We saw no significant effect of IL-21 alone on Blimp-1 or T-bet expression in MAIT cells, but did observe significant upregulation of pSTAT1, pSTAT3, and pSTAT5, with IL-21 having the strongest effect on pSTAT3. This was expected as IL-21 predominantly signals through STAT3.(38) In CD4^+^ T cells, increased STAT3 signalling has been associated with upregulation of GrzB and decreased expression of IL-2, IFN-γ, and TNF-α.(59) Therefore, it is possible that enhanced STAT3 signalling contributes to IL-21-driven changes in MAIT cell GrzB and cytokine expression. Supporting this, we observed minimal impact of IL-21 on IFN-γ production by MAIT cells, alongside significantly less IL-2 and TNF-α in cell culture supernatants when IL-21 was added to αCD3/CD28-stimulated Vα7.2^+^ cells.

However, it is unlikely that STAT3 is the sole driver of IL-21-induced GrzB production in MAIT cells. A previous study reported that in individuals with loss-of-function mutations in STAT3, whilst they had reduced frequency of MAIT cells, MAIT cells still retained normal GrzB production.(41) While our results highlight that IL-21 facilitates activation of multiple STAT proteins in MAIT cells, the precise mechanism for how IL-21 modulates MAIT cell cytotoxic function at the transcriptional level remains elusive.

The effect of IL-21 on MAIT cell function has not been extensively studied. However, a recent report observed that in a 5-OP-RU-treated THP-1/ Vα7.2^+^ 24-hr co-culture, IL-21 preserved MAIT cell GrzB content and significantly upregulated IL-10 production but did not impact TNF-α, IFN-γ, or IL-17 expression.(60) This aligns with our observations that IL-21 does not induce MAIT cell pro-inflammatory cytokine profile, however, in contrast to their findings, we did not observe any impact of IL-21 on IL-10 production by MAIT cells in our LEGENDplex™ analysis. This could be due to the difference in stimulus (5-OP-RU-treated THP-1 vs αCD3/CD28 beads used in our study) and timing (24 hrs vs 6 hrs in our study). We also observed a less pronounced upregulation of GrzB by MAIT cells from isolated Vα7.2^+^ cell cultures stimulated with IL-21 and αCD3/CD28 compared to MAIT cells in 5-OP-RU/IL-21-treated PBMC cultures. Furthermore, in contrast to the present study, Kurioka et al.(19) found no enhancement of GrzB on MAIT cells following 24 hr stimulation with IL-21 and observed a non-significant increase in perforin expression. We also observed a modest increase in perforin expression but a significant upregulation of GrzB after 24 hrs of IL-21 treatment. Differences in cell types (enriched CD8^+^ T cells vs PBMCs) may explain these discrepancies. IL-21 may induce production of cytokines by other cells present in PBMCs which may affect MAIT cell GrzB and perforin production. Thus, the *in vitro* culture system used in this study is a limitation for assessing MAIT cell responses to IL-21.

Additionally, blocking IL-21R during 6 hr treatment of PBMCs with *E. coli* significantly reduced MAIT cell GrzB and perforin expression, however, inhibiting MR1 and T1-IFNs had a greater impact on MAIT cell response, confirming that TCR signalling and T1-IFNs are the key signals driving early MAIT cell response to *E. coli.*(29) While we saw modest production of IL-21 by PB MAIT and CD4^+^ T cells at this time point, IL-21 is unlikely to be the primary signal regulating MAIT cell cytotoxicity against *E. coli*, as its effect was comparable to sub-optimal concentrations of T1-IFNs.

Previous studies primarily focus on MAIT cells as producers of IL-21 that support B cell function.(34-37, 61) Our results highlight a potential autocrine role for IL-21 in MAIT cells. Autocrine IL-21 production has been reported in Th17 and CD4^+^ T cells,(62-64) and IL-27-induced IL-21 modulates GrzB production by CD8^+^ T cells in an autocrine manner.(44) Given IL-21 induces gene expression of its own receptor,(65) it is plausible that MAIT cell derived IL-21 may modulate MAIT cell cytotoxicity via an autocrine mechanism. However, it is unclear whether tissue-resident MAIT cells respond to IL-21, as the present study used PBMC-derived MAIT cells which produce minimal IL-21.(34, 37) Further research is needed to explore the impact of IL-21 on tissue MAIT cells and to determine whether MAIT cell-derived IL-21 enhances MAIT cell cytotoxicity *in vivo*.

In conclusion, we have demonstrated a previously unexplored role for IL-21 in enhancing MAIT cell cytotoxic potential. The ability for IL-21 to selectively augment MAIT cell cytotoxic functions highlights a potential mechanism for modulating MAIT cell cytotoxic function independently of MAIT cell activation and cytokine production. Furthermore, our findings raise the possibility that MAIT cells may regulate their own cytotoxic function through IL-21. These findings have implications for the development of IL-21 and MAIT cell directed therapies and inform further research into how IL-21 impacts MAIT cell function in different contexts.

## Acknowledgements

This work was supported by the Health Research Council of New Zealand (J.E.U.) and Department Professor Sandy Smith Memorial Scholarship (L.E.W).

## Author contributions

L. E.W., R.L performed the experiments. L.W analysed the data. L.W., J.E.U. designed the experiments. J.E.U. managed the study. S.G and A.V. synthesised the 5-A-RU. L.E.W., J.E.U. conceived the work and wrote the manuscript which was revised and approved by all authors.

## Ethics approval

Ethics approval was obtained from the University of Otago Human Ethics Committee (Health): H14/046.

## Abbreviations

TCR: T cell receptor
5-OP-RU: 5-(2-oxopropylideneamino)-6-d-ribitylaminouracil
MAIT cell: mucosal associated invariant T cells
PBMC: peripheral blood mononuclear cells
IL-21: interleukin-21
GrzB: granzyme B

