## Supplementary figures 1-8 for "IL-21 selectively augments cytotoxic potential of antigen-activated MAIT cells"

#### Figure Legends:

**Supporting information Fig 1: Gating strategy for gating MAIT and B cells from PBMC**

Gating strategy for gating MAIT cells ($CD3^{+}CD8^{+}CD161^{++}$ $V\alpha 7.2^{+}$) and B cells ($CD3^{-}CD19^{+}$) from PBMCs for assessing IL-21R expression in Fig. 1.

**Supporting information Fig 2: Gating strategy for gating MAIT cells from PBMC for assessing expression of cytokines and cytotoxic molecules**

(A) Gating strategy for gating MAIT cells from PBMCs; used in Fig. 2 and Fig. 4-6. (B) PBMCs were activated with *E. coli* for 6 hr; expression of cytotoxic molecules or cytokines on unstimulated (black) or *E. coli*-stimulated (red) MAIT cells are shown as flow plots. Gates for GrzA, GrzB, TNF-α, and IFN-γ were set using untreated samples. Percentage positive cells for each gate in untreated (black) and *E. coli*-treated (red) are also shown. In general, MAIT cells were gated as $CD3^{+}CD8^{+}CD161^{++}V\alpha 7.2^{+}$. In some experiments (Fig. 5), MAIT cells were gated as $CD3^{+}CD161^{++}V\alpha 7.2^{+}$.

**Supporting information Fig 3: Gating strategy for assessing purity of isolated $V\alpha 7.2^{+}$ cells**

Gating strategy for gating MAIT cells from isolated $V\alpha 7.2^{+}$ cells in Fig. 3 and for assessing purity of MACS isolation. A representative graph showing %$V\alpha 7.2^{+}$ of live and %MAIT cells of $V\alpha 7.2^{+}$ from two independent experiments is also shown (n=6).

**Supporting information Fig 4: Gating strategy for gating $CD4^{+}$ T cells and MAIT cells from PBMCs**

Gating strategy for gating $CD4^{+}$ T cells and MAIT cells from PBMCs for analysis of IL-21 expression in Fig. 6H-J.

**Supporting information Fig 5: Gating strategy for cytotoxic assay**

Gating strategy for cytotoxic assay. Cells were first gated based on size and granularity using BCL only and Vα7.2$^+$ only controls to set gates. Cells were then gated as single cells before gating for live cells. For cytotoxic assays using PBMCs as effector cells, MAIT cells were first gated as CD3$^+$ before gating for MAIT cells (CD161$^{++}$Vα7.2$^+$).

**Supporting information Fig 6: IL-21 does not enhance cytokine production or granulysin expression by MAIT cells**

(A) PBMCs were treated with 5-OP-RU with or without IL-21 for 6 hrs; brefeldin A was added for the final 4 hrs. Granulysin MFI and the frequency of MAIT cells expressing IFN-γ and TNF-α were assessed by flow cytometry. (B-C) PBMC were treated with 5-OP-RU with or without IL-21 or PMA/ionomycin for 6 hrs; brefeldin A was added for the final 4 hrs. The percentage of IL-17A$^+$ MAIT cells was assessed by flow cytometry (B). (C) Representative flow plots for each treatment are shown; gates were set using untreated samples. Each biological replicate (n=8 (A), n=7 (B)) and the mean ± SEM from two independent experiments are shown. Data were analysed with Friedman's test with Dunn's multiple comparison post-test or repeated measures one-way ANOVA with Sidak's multiple comparison post-test. MFI = median fluorescence intensity. ns = no significance, * ($p<0.05$),** ($p<0.01$).

**Supporting information Fig 7: Time of exposure to IL-21 influences expression of cytotoxic molecules**

(A-D) PBMCs were treated with IL-21 for 6, 12, 24, 48 or 72 hrs; 0 hrs represents untreated. Brefeldin A was added for the final 4 hrs of each incubation. MFI of perforin (A) and GrzA (C), and granulysin (D) and the frequency of MAIT cells expressing GrzB (B) were assessed by flow cytometry. Each data point represents the mean ± SEM (n=9-10) from two independent experiments. Samples 0-24 hours were stained as one group and 48-72 hr were stained as one group. MFI = median fluorescence intensity.

**Supporting information Fig 8: IL-21 enhances MAIT cell responses to 5-OP-RU in combination with other cytokines**

(A-D) PBMCs were treated with 5-OP-RU and combinations of IL-21, IFN-α, and IFN-β for 6 hrs. (E-H) PBMCs were treated with combinations of IL-2, IL-7, IL-15, and IL-21, with or without 5-OP-RU for 6 hrs. Brefeldin A was added for the final 4 hrs. Perforin MFI (A, C, E, G) and the frequency of MAIT cells expressing GrzB (B, D, F, H) were assessed by flow cytometry. Each biological replicate (n=8 (A-B), n=7 (C-D), n=6, (E-H)) and the mean ± SEM from two independent experiments are shown. Data were analysed with Friedman's test with Dunn's multiple comparison post-test or repeated measures one-way ANOVA with Sidak's multiple comparison post-test. MFI = median fluorescence intensity. ns = no significance, *$p<0.05$,**$p<0.01$,***$p<0.001$,****$p<0.0001$.

### FIGURE S1

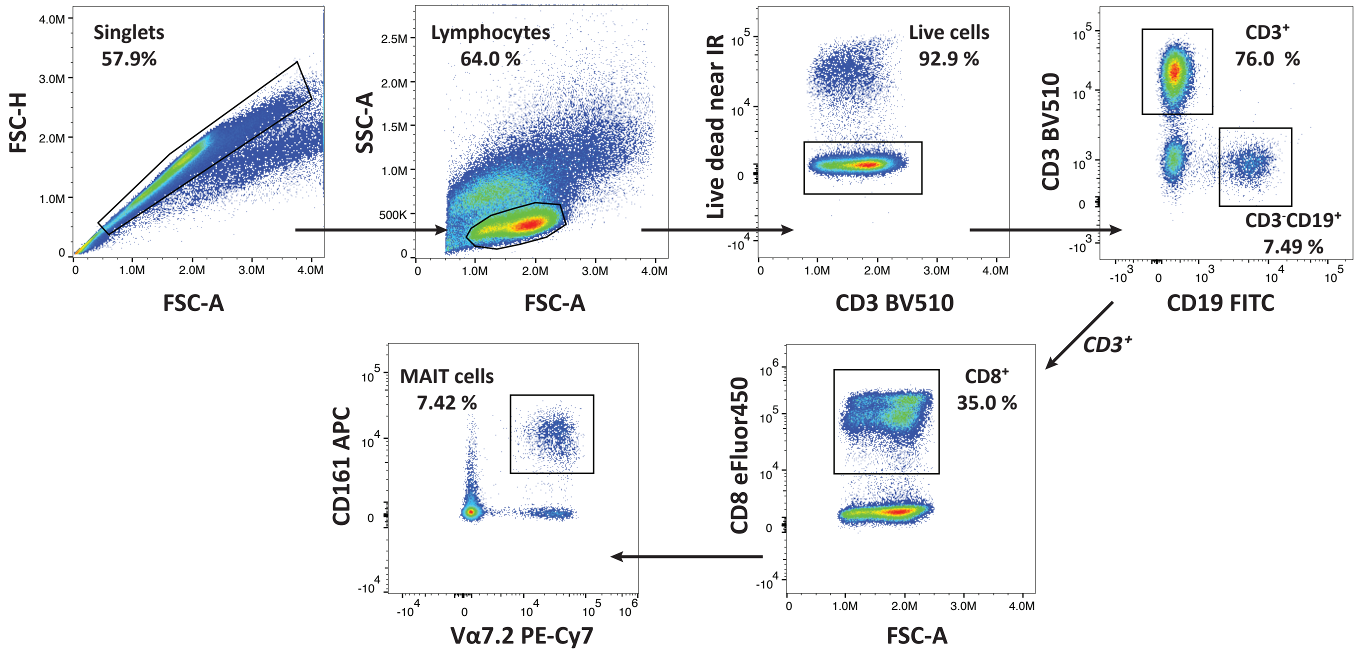
Singlets
57.9%
FSC-H
FSC-A
Lymphocytes
64.0 %
SSC-A
FSC-A
Live cells
92.9 %
Live dead near IR
CD3 BV510
CD3+
76.0 %
CD3-CD19+
7.49 %
CD3 BV510
CD19 FITC
CD3+
CD8+
35.0 %
CD8 eFluor450
FSC-A
MAIT cells
7.42 %
CD161 APC
Vα7.2 PE-Cy7

### FIGURE S2

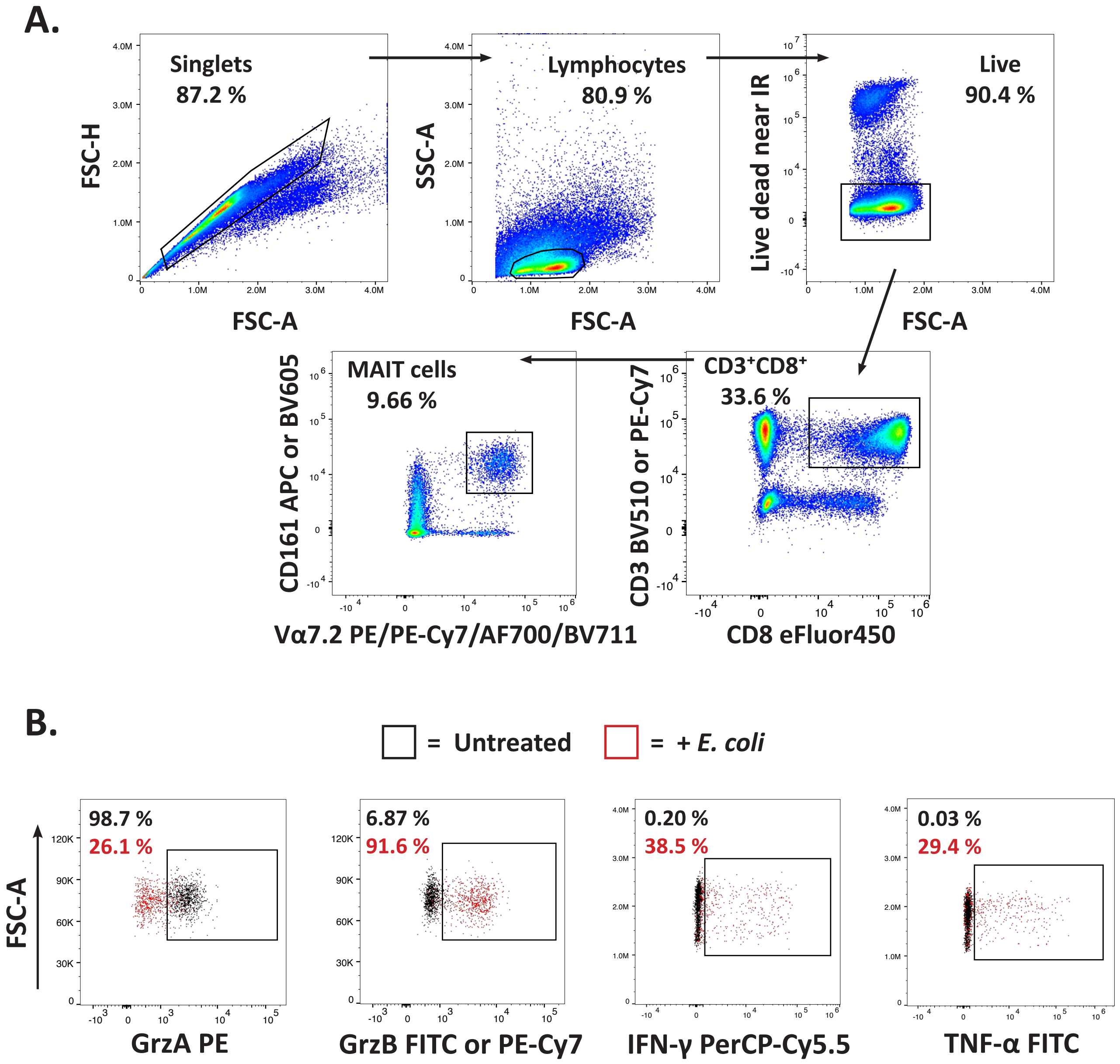
A.
Singlets
87.2 %
FSC-H
FSC-A
Lymphocytes
80.9 %
SSC-A
FSC-A
Live
90.4 %
Live dead near IR
FSC-A
CD3+CD8+
33.6 %
CD3 BV510 or PE-Cy7
CD8 eFluor450
MAIT cells
9.66 %
CD161 APC or BV605
Vα7.2 PE/PE-Cy7/AF700/BV711
B.
= Untreated
= + E. coli
FSC-A
98.7 %
26.1 %
GrzA PE
6.87 %
91.6 %
GrzB FITC or PE-Cy7
0.20 %
38.5 %
IFN-γ PerCP-Cy5.5
0.03 %
29.4 %
TNF-α FITC

### FIGURE S3

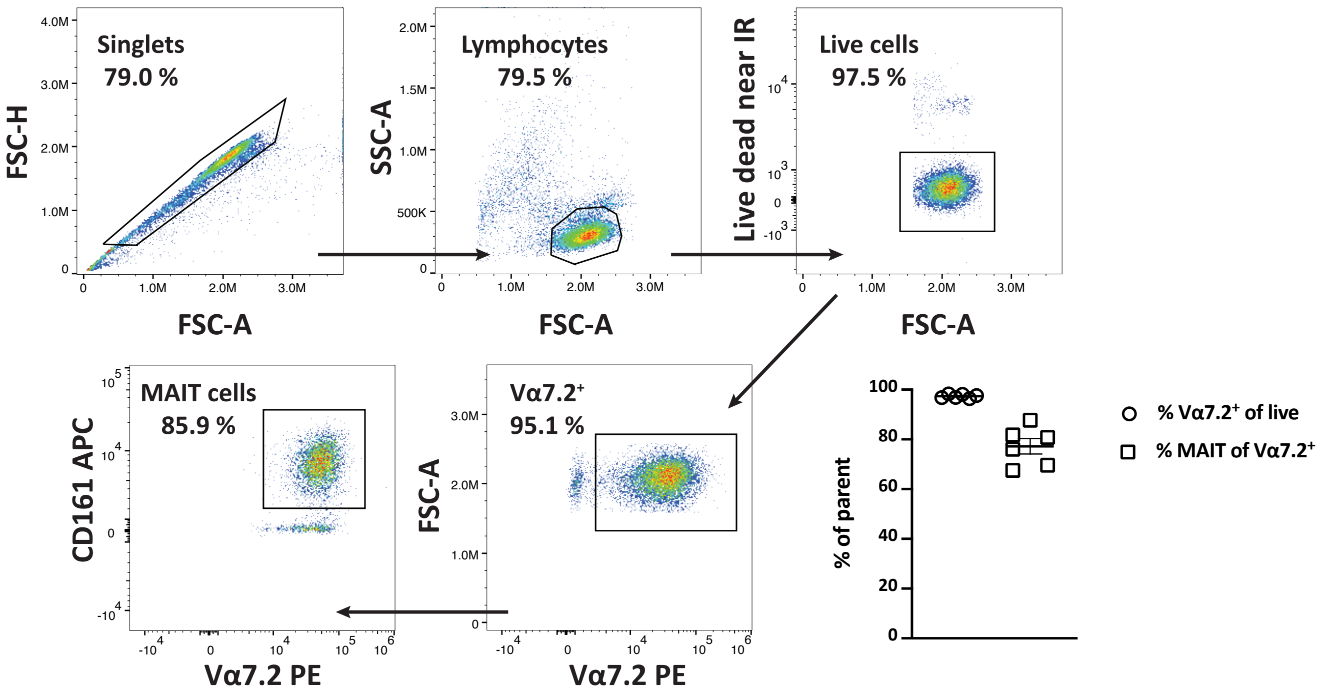
Singlets
79.0 %
FSC-H
FSC-A
Lymphocytes
79.5 %
SSC-A
FSC-A
Live cells
97.5 %
Live dead near IR
FSC-A
MAIT cells
85.9 %
CD161 APC
Vα7.2 PE
Vα7.2⁺
95.1 %
FSC-A
Vα7.2 PE
% of parent
% Vα7.2⁺ of live
% MAIT of Vα7.2⁺

### FIGURE S4

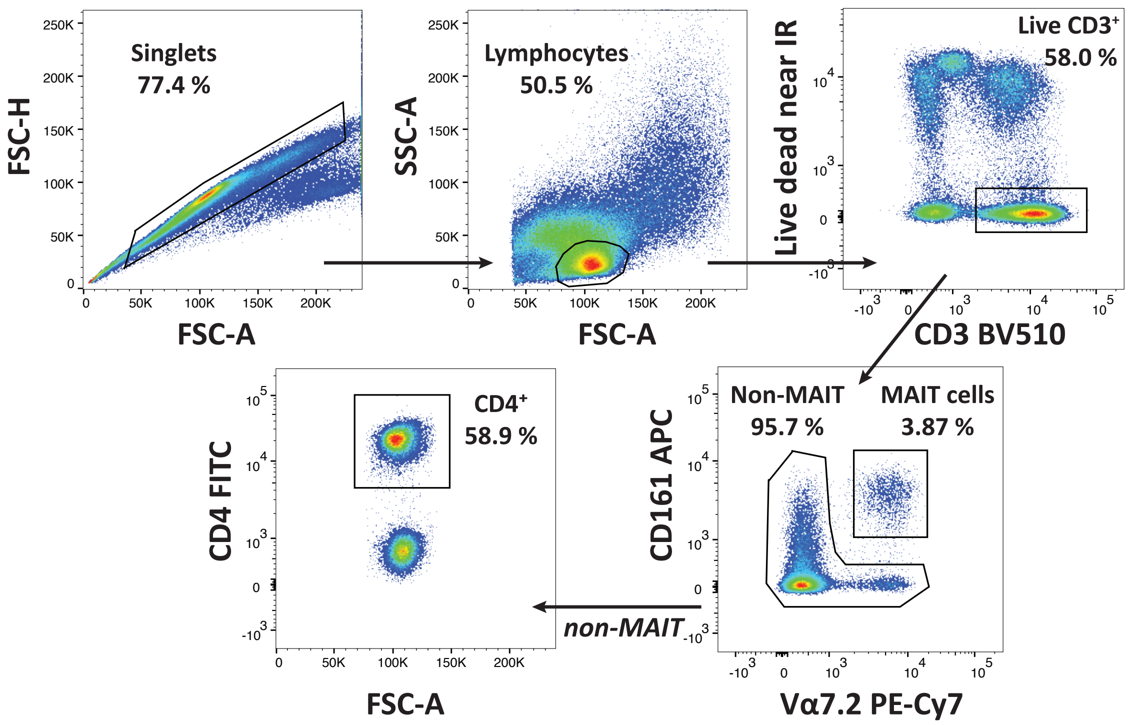

Singlets
77.4 %
FSC-H
FSC-A
Lymphocytes
50.5 %
SSC-A
FSC-A
Live CD3+
58.0 %
Live dead near IR
CD3 BV510
Non-MAIT
95.7 %
MAIT cells
3.87 %
CD161 APC
Vα7.2 PE-Cy7
non-MAIT
CD4+
58.9 %
CD4 FITC
FSC-A

### FIGURE S5

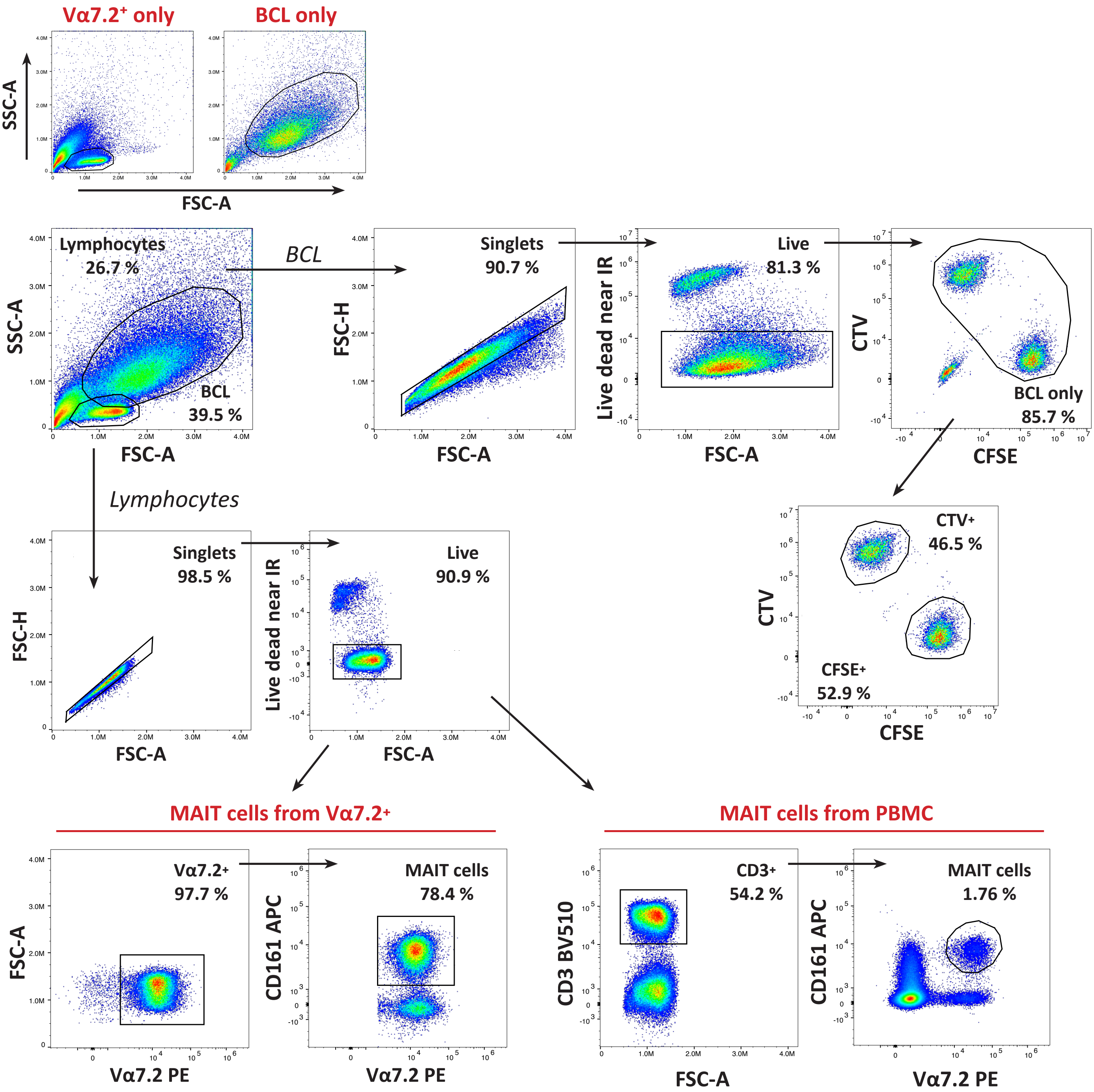

Vα7.2+ only
BCL only
SSC-A
FSC-A
Lymphocytes
26.7 %
BCL
39.5 %
BCL
Singlets
90.7 %
FSC-H
Live dead near IR
Live
81.3 %
CTV
CFSE
BCL only
85.7 %
Lymphocytes
Singlets
98.5 %
Live
90.9 %
CTV+
46.5 %
CFSE+
52.9 %
MAIT cells from Vα7.2+
MAIT cells from PBMC
Vα7.2+
97.7 %
Vα7.2 PE
CD161 APC
MAIT cells
78.4 %
CD3 BV510
CD3+
54.2 %
MAIT cells
1.76 %

### FIGURE S6

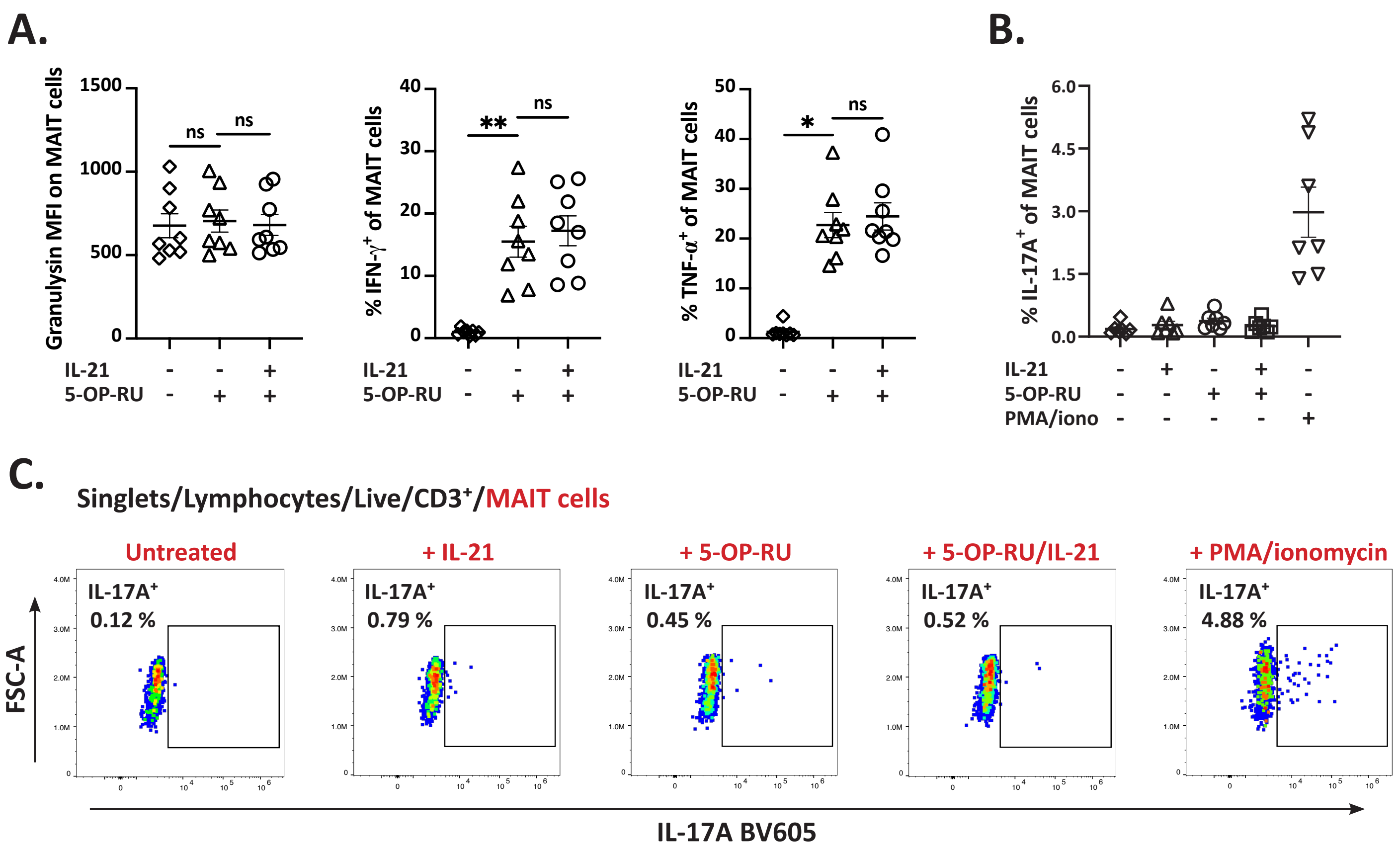

A.
Granulysin MFI on MAIT cells
1500
1000
500
0
ns
ns
IL-21 - - +
5-OP-RU - + +
% IFN-γ+ of MAIT cells
40
30
20
10
0
**
ns
IL-21 - - +
5-OP-RU - + +
% TNF-α+ of MAIT cells
50
40
30
20
10
0
*
ns
IL-21 - - +
5-OP-RU - + +
B.
% IL-17A+ of MAIT cells
6.0
4.5
3.0
1.5
0.0
IL-21 - + - + -
5-OP-RU - - + + -
PMA/iono - - - - +
C.
Singlets/Lymphocytes/Live/CD3+/MAIT cells
Untreated
IL-17A+ 0.12 %
+ IL-21
IL-17A+ 0.79 %
+ 5-OP-RU
IL-17A+ 0.45 %
+ 5-OP-RU/IL-21
IL-17A+ 0.52 %
+ PMA/ionomycin
IL-17A+ 4.88 %
FSC-A
IL-17A BV605

#### FIGURE S7

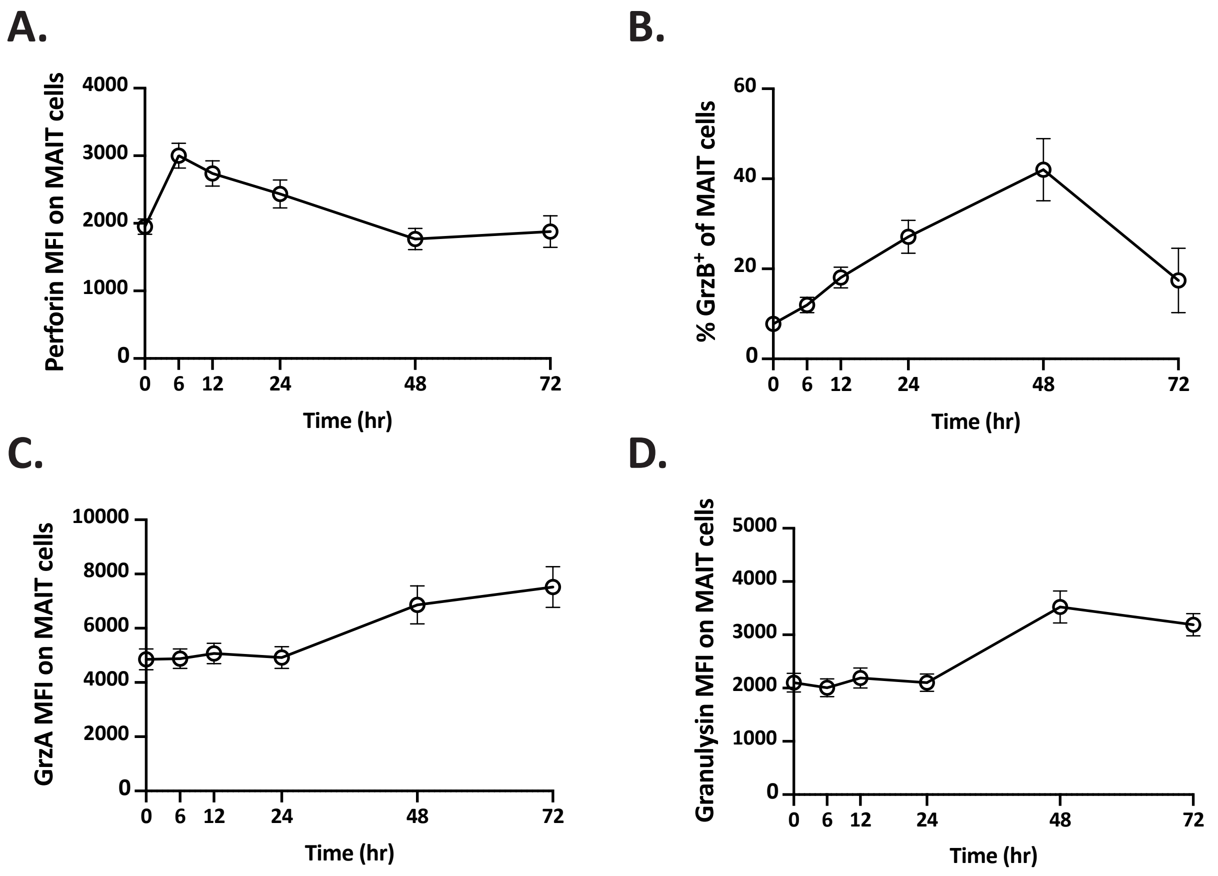
A.
Perforin MFI on MAIT cells
4000
3000
2000
1000
0
0 6 12 24 48 72
Time (hr)
B.
% GrzB+ of MAIT cells
60
40
20
0
0 6 12 24 48 72
Time (hr)
C.
GrzA MFI on MAIT cells
10000
8000
6000
4000
2000
0
0 6 12 24 48 72
Time (hr)
D.
Granulysin MFI on MAIT cells
5000
4000
3000
2000
1000
0
0 6 12 24 48 72
Time (hr)

### FIGURE S8

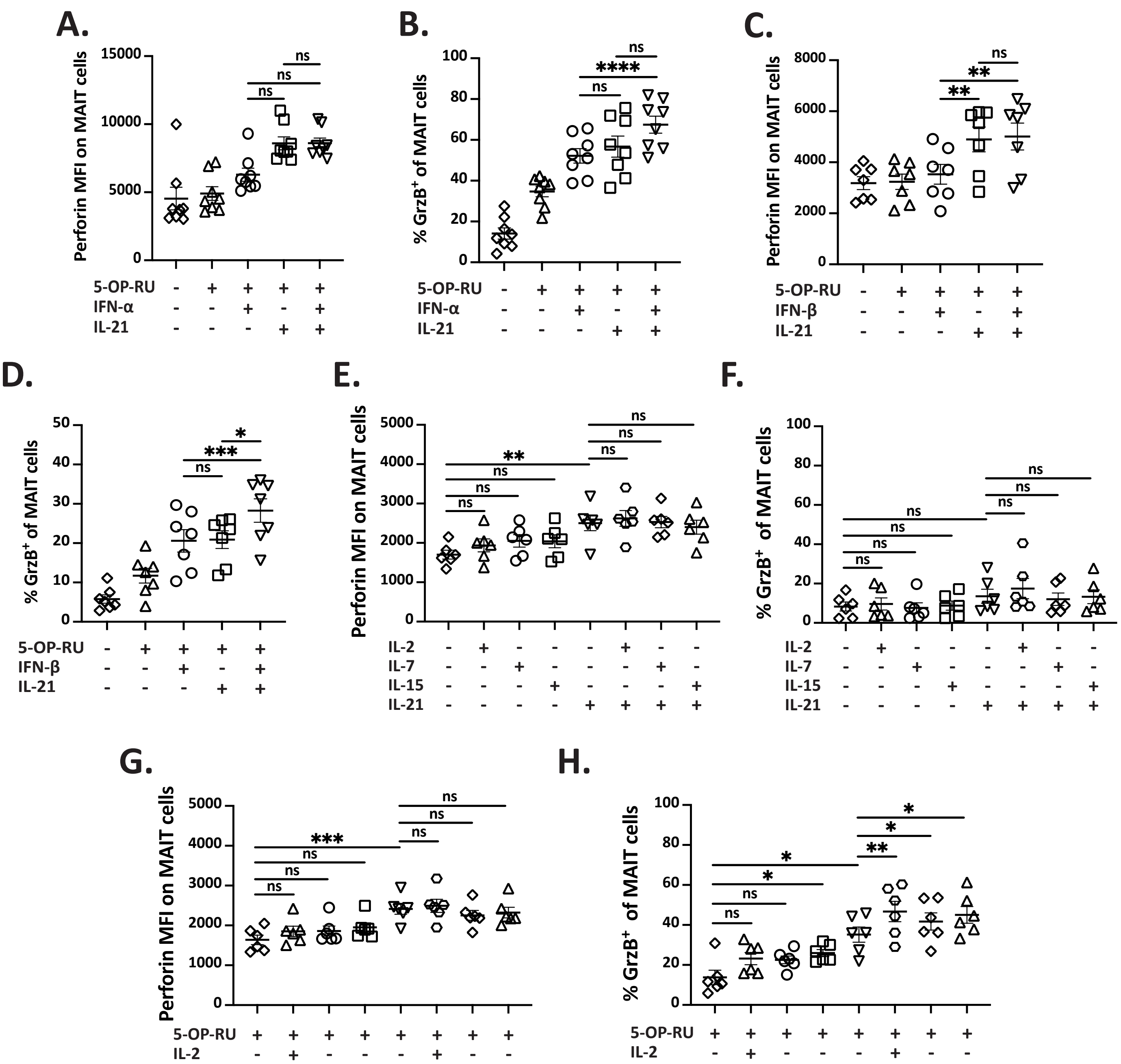
A.
Perforin MFI on MAIT cells
5-OP-RU
IFN-α
IL-21
B.
% GrzB+ of MAIT cells
5-OP-RU
IFN-α
IL-21
C.
Perforin MFI on MAIT cells
5-OP-RU
IFN-β
IL-21
D.
% GrzB+ of MAIT cells
5-OP-RU
IFN-β
IL-21
E.
Perforin MFI on MAIT cells
IL-2
IL-7
IL-15
IL-21
F.
% GrzB+ of MAIT cells
IL-2
IL-7
IL-15
IL-21
G.
Perforin MFI on MAIT cells
5-OP-RU
IL-2
IL-7
IL-15
IL-21
H.
% GrzB+ of MAIT cells
5-OP-RU
IL-2
IL-7
IL-15
IL-21
